# The NAD^+^ Precursor Nicotinamide Riboside Rescues Mitochondrial Defects and Neuronal Loss in iPSC derived Cortical Organoid of Alpers’ Disease

**DOI:** 10.1101/2023.07.02.547346

**Authors:** Yu Hong, Zhuoyuan Zhang, Tsering Yangzom, Anbin Chen, Bjørn Christian Lundberg, Evandro Fei Fang, Gareth John Sullivan, Charalampos Tzoulis, Laurence A. Bindoff, Kristina Xiao Liang

**Author notes:** **Correspondence to:** Kristina Xiao Liang, Department of Clinical Medicine (K1), University of Bergen, Jonas Lies vei 87, P. O. Box 7804, 5021 Bergen, Norway. These authors contributed equally to this work as first authorship (Yu Hong, Zhuoyuan Zhang).

## Abstract

Alpers’ syndrome is an early-onset neurodegenerative disorder usually caused by biallelic pathogenic variants in the gene encoding the catalytic subunit of polymerase-gamma (POLG), which is essential for mitochondrial DNA (mtDNA) replication. The disease is progressive, incurable, and inevitably it leads to death from drug-resistant status epilepticus. The neurological features of Alpers’ syndrome are intractable epilepsy and developmental regression, with no effective treatment; the underlying mechanisms are still elusive, partially due to lack of good experimental models. Here, we generated the patient-derived induced pluripotent stem cells from one Alpers’ patient carrying the compound heterozygous mutations of A467T (c.1399G>A) and P589L (c.1766C>T), and further differentiated them into cortical organoids and neural stem cells (NSCs) for mechanistic studies of neural dysfunction in Alpers’ syndrome. Patient cortical organoids exhibited a phenotype that faithfully replicated the molecular changes found in patient postmortem brain tissue, as evidenced by cortical neuronal loss and depletion of mtDNA and complex I (CI). Patient NSCs showed mitochondrial dysfunction leading to ROS overproduction and downregulation of the NADH pathway. More importantly, the NAD^+^ precursor nicotinamide riboside (NR) significantly ameliorated mitochondrial defects in patient brain organoids. Our findings demonstrate that the iPSC model and brain organoids are good *in vitro* models of Alpers’ disease; this first-in-its-kind stem cell platform for Alpers’ syndrome enables therapeutic exploration and has identified NR as a viable drug candidate for Alpers’ disease and, potentially, other mitochondrial diseases with similar causes.

## Introduction

Alpers’ syndrome, a devastating early-onset neurodegenerative disorder is tightly linked to abnormalities in the function of mitochondria and mutations in the catalytic subunit of polymerase gamma (*POLG*) gene [1]. The syndrome, which typically follows an autosomal mode of inheritance, manifests a range of debilitating symptoms including intractable seizures, developmental regression, hypotonia, ataxia, and liver failure. In some severe instances, it can lead to premature death [2]. Symptoms usually emerge between first and third years of life, with the disorder estimated to affect 1 in 100,000 newborns [3].

Unfortunately, Alpers’ syndrome’s pathogenesis remains obscure, making the development of direct treatment modalities challenging. Consequently, current interventions focus on symptomatic relief and providing supportive care. Certain drugs previously utilized, like valproic acid, are now largely discarded due to their severe hepatotoxic effects and persistence of drug-resistant epilepsy [4, 5]. This underlines the crucial need for continuing research into the root cause of Alpers’ syndrome, which would pave the way for devising strategies to limit disease progression and generate efficacious treatments.

The *POLG* gene, situated on the long arm of chromosome 15, encodes for polymerase γ (Pol γ), an enzyme chiefly responsible for replication and repair of mitochondrial DNA (mtDNA) [1]. More than 90% of Alpers’ syndrome cases are linked to mutations in the *POLG* gene. These mutations are typically inherited in an autosomal recessive manner, often from parents who are asymptomatic carriers. The mutations commonly observed include homozygous p.A467T or p.W748S transitions and heterozygous p.T851A and p.R1047W mutations, all of which impact the function and phenotypic manifestation of Pol γ A gradual deficiency in Pol γ contributes directly to mtDNA depletion, instigating serious harm to the mitochondrial respiratory chain, particularly leading to the breakdown of CI [6, 7]. The ensuing severe mitochondrial dysfunction manifests as ataxia, hepatic failure, and even premature death. Since Alpers’ syndrome is a metabolic disorder, deepening our understanding of its pathogenesis could pave the way for a more comprehensive grasp of mitochondrial metabolism. This could potentially open new avenues for more effective treatments.

Unraveling the pathogenesis of neurodegenerative diseases, including Alpers’ disease, is a complex task due largely to the intricacy of the human nervous system. To date, most research efforts have relied on genetically modified cell lines that is limited to relatively single immortalized cell lines [8] or rodent primary neural stem cells (NSCs) [9]. However, these models may not adequately represent the multifaceted in vivo microenvironment of the human body. *In vivo* studies using animal models provide a simplified, cost-effective approach to neurodegeneration research. . Several mouse models have been developed to recapitulate key features of Alpers’ disease, enabling the study of disease progression, mitochondrial dysfunction, and associated phenotypic changes. These models often involve the targeted disruption or manipulation of genes involved in mitochondrial function, such as POLG [10] and Twinkle [11]. By selectively introducing mutations or altering gene expression in these models, researchers can investigate the specific effects on mitochondrial DNA replication, energy metabolism, and oxidative stress, all of which are implicated in the pathogenesis of Alpers’ disease. Nevertheless, their applicability is constrained due to inherent differences between human and animal models [12]. Furthermore, neuroprotective drugs showing promise in animal models often disappoint when tested in human clinical trials [13]. Given these limitations, there’s an urgent need for a more accurate disease model that truly emulates the complexities of neurodegenerative diseases in humans. Such a model could unlock insights into disease mechanisms and expedite the development of effective therapeutic strategies.

Introduced by Shinya Yamanaka in 2007, human induced pluripotent stem cells (iPSCs), have emerged as a promising source of cells for in vitro studies and clinical applications in humans and patients [14]. This innovative approach reverts human somatic cells, such as skin fibroblasts or peripheral blood mononuclear cells, back to a pluripotent state using various combinations of transcription factors [15]. iPSCs derived from specific diseases can retain the patient’s individual genetic mutations and avoid potential interspecies variation, offering a tremendous to study neurodegenerative diseases at the cellular level [16]. To date, patient-specific iPSCs have been been extensively utilized in the creation of diverse neuronal models [17] and in the exploration of mitochondrial diseases caused by mtDNA mutations, such as Mitochondrial Encephalomyopathy, Lactic Acidosis, and Stroke-like Episodes (MELAS) syndrome [18, 19], Myoclonic Epilepsy with Ragged-Red Fibers (MERRF) syndrome [20], Pearson syndrome (PS) [21] and Hypertrophic Cardiomyopathy (HCM) [22]. However, most cellular assays primarily rely on traditional two-dimensional (2D) culture systems, which are increasingly recognized as inadequate for mimicking complex natural environments. Recent evidence supports the idea that three-dimensional (3D) cell culture systems, such as brain organoids derived from iPSCs, could address this limitation effectively. These 3D systems accurately reflect *in vivo* microenvironment [23, 24], providing robust models to emulate the disease profiles of neurodegenerative disease and conduct drug testing.

Nicotinamide adenine dinucleotide in its oxidized form (NAD^+^) is a vital molecule for life and health, playing a particularly crucial role in neuroprotection. NAD^+^’s fundamental molecular functions extend beyond its roles in energy metabolism and production, including glycolysis, β-oxidation, the TCA cycle, and oxidative phosphorylation (OXPHOS). It also contributes to cellular repair and resilience, factors that underpin its ability to protect the brain [25, 26]. Research conducted on postmortem brain tissues, iPSC-derived cells, and animal models indicates a decline in NAD^+^ levels in the aging brain, as well as in prevalent neurodegenerative diseases, such as Alzheimer’s disease (AD), Parkinson’s disease (PD), Amyotrophic lateral sclerosis (ALS), and Huntington’s disease (HTT) [27]. The belief that the reduction in NAD^+^ is a primary cause, rather than a simple correlate, is bolstered by evidence showing that NAD^+^ augmentation-achieved through supplementation with NAD^+^ precursors like as nicotinamide riboside (NR) and nicotinamide mononucleotide (NMN), can alleviate disease pathologies in laboratory models of these diseases [27]. Notably, clinical studies have demonstrated that NR is not only safe (up to 2 g/day for up to 3 months) and bioavailable, but it can also ameliorate some syndromes in patients with PD [28] and ALS [29].

To date, there has been no reported use of the iPSCs model for the study of Alpers’ syndrome resulting from *POLG* mutation. In this study, we established an iPSCs model of Alpers’ syndrome, generated from the skin fibroblasts of affected patients. These cells were further differentiated into 2D NSCs and 3D cortical to investigate the varied manifestations and pathogenesis of Alpers’ disease. Both the iPSCs and NSCs derived from Alpers’ patients exhibited some degree of mitochondrial dysfunction, the dysfunction being more pronounced in NSCs.

With the use of a cortical organoid model derived from iPSCs, we found that the mitochondrial alterations in the cortical organoids from Alpers’ patients mirrored those observed in 2D NSCs. Additionally, a transcriptomic analysis revealed a downregulation of the NADH pathway and reduced expression of mitochondrial transcripts in both reprogrammed iPSCs and induced NSCs. This provided key insights into potential therapeutic approaches. Notably, subsequent studies in the cortical organoid model demonstrated that nicotinamide riboside (NR) significantly rectified the structural disruptions and mitochondrial defects in brain organoids generated from an Alpers’ syndrome patient. For the first time, we have constructed a stem cell model for Alpers’ syndrome, paving the way for precise mechanistic studies and the exploration of potential drug development for this currently incurable disease.

With the use of a cortical organoid model derived from iPSCs, we found that the mitochondrial alterations in the cortical organoids from Alpers’ patients mirrored those observed in 2D NSCs. Additionally, a transcriptomic analysis revealed a downregulation of the NADH pathway and reduced expression levels of mitochondrial transcripts in both reprogrammed iPSCs and induced NSCs. provided key insights into potential therapeutic approaches. Notably, subsequent studies in the cortical organoid model demonstrated that NR significantly rectified the structural disturbances and mitochondrial defects in brain organoids generated from an Alpers’ syndrome patient. For the first, we have constructed a stem cell model for Alpers’ syndrome, paving the way for precise mechanistic studies and the exploration of potential drug development for this currently incurable disease.

## Results

### Alpers’s iPSCs manifested a mild impairment of mitochondrial function compared to controls

We generated iPSCs from a patient with Alpers’ syndrome who carries heterozygous mutations A467T (c.1399G>A) and P589L (c.1766C>T) (Figure 1A). Both mutations are located in the linker region that is in close proximity to the auxiliary subunit (Figure 1B). To validate the pluripotency and cellular purity of the iPSCs, we utilized flow cytometry, which confirmed that over 97% of the iPSCs derived from Alpers’ patients expressed the pluripotent marker NANOG (Supplemental Figure 1). Fluorescent staining further confirmed that the iPSC colonies expressed the pluripotent markers OCT4 and SSEA4 (Figure 1C). Sanger sequencing analysis corroborated that two mutations present in the iPSCs were consistent with those identified in the patient (Figure 1D). We utilized two age and/or gender-matched control lines, derived from control Detroit 551 fibroblasts (ATCC CCL 110TM) and CRL2097 fibroblasts (ATCC CRL-2097™), as disease-free controls. We observed that the iPSCs derived from Alpers’ patients exhibited typical colony morphology, similar to the control iPSCs (Supplemental Figure 2).

**Figure 1.**
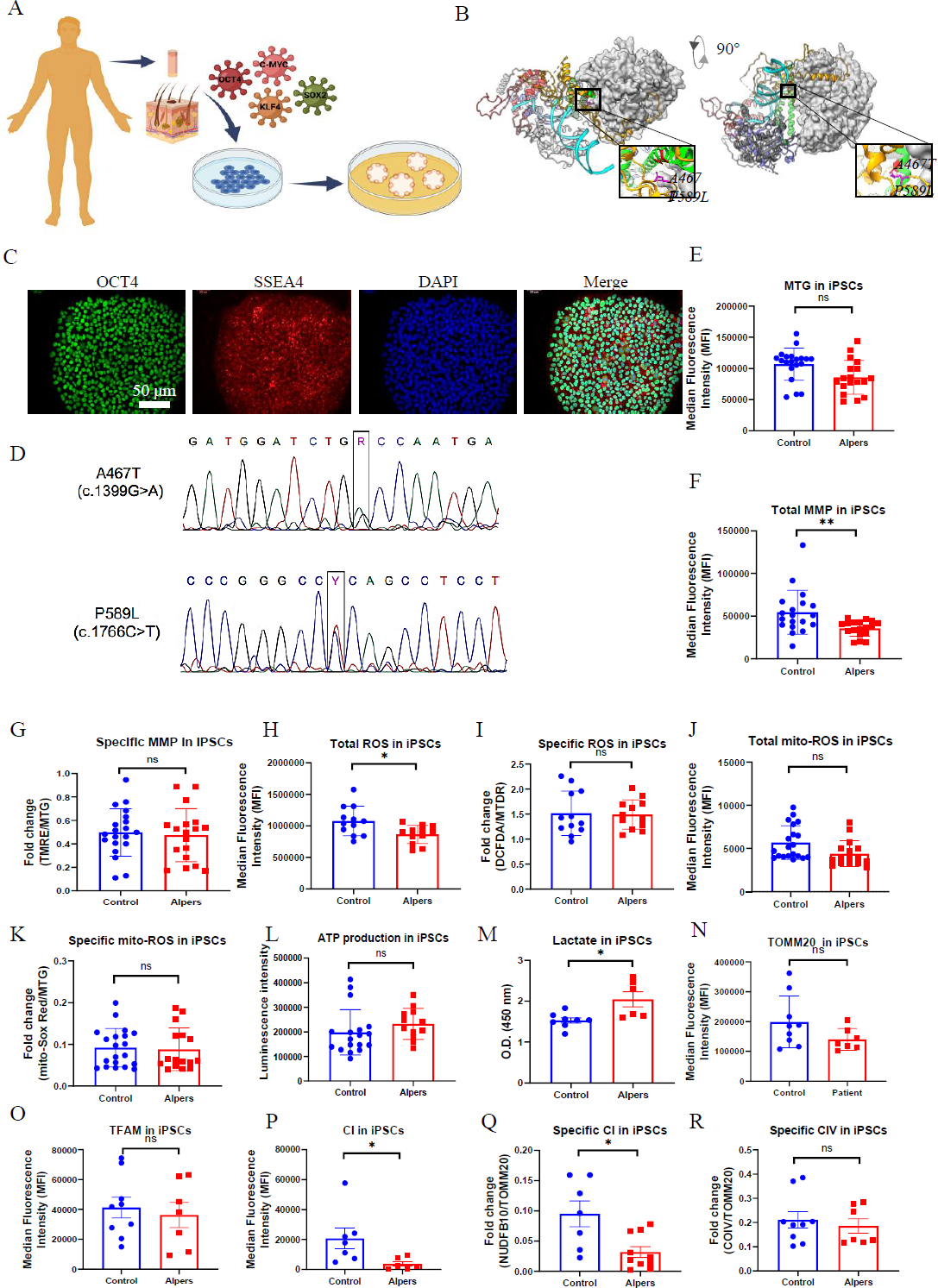
Measurement of mitochondrial function, mtDNA alteration and mitochondrial complexes in Alpers’ iPSCs. A. Illustration of reprogramming of Alpers’ patients’ fibroblasts into iPSCs. B. Molecular Structure of POLG protein and position of A467T and P589L mutations. C. Representative fluorescent images of Alpers’ iPSCs for pluripotent markers OCT4 and SSEA4. Nuclei are stained with DAPI (blue). Scale bar is 50 µm. D. A467T and P589L mutation verified by sanger sequencing using Alpers’ reprogrammed iPSCs. E. Flow cytometric analysis for mitochondrial volume using MTG in iPSCs. F-G. Flow cytometric analysis for total MMP and specific MMP in iPSCs. H-K. Flow cytometric analysis for total ROS, specific ROS, total mito-ROS, and specific mito-ROS in iPSCs. L-M. Flow cytometric analysis for ATP and lactate production in iPSCs. N. Flow cytometric analysis for TOMM20 levels in iPSCs. O. Flow cytometric analysis for TFAM in iPSCs. P-R. Flow cytometric analysis for CI, specific CI, CIV, specific CIV levels in iPSCs.

To determine if there were changes in mitochondrial function in iPSCs reprogrammed from Alpers’ patient fibroblasts, we assessed mitochondrial membrane potential (MMP), mitochondrial mass, ROS levels and intracellular energy production. Mitochondrial mass and MMP were measured. Each was measured by flow cytometry after double-staining cells with MitoTracker Green (MTG) and tetramethylrhodamine ethyl ester (TMRE). To gauge MMP levels per mitochondrion, we measured the ratio between total MMPs and MTG, providing relative measurements of specific MMPs.

Our results showed that mitochondrial volume measured by MTG (Figure 1E) and total MMPs assessed by TMRE (Figure 1F) were significantly reduced in Alpers’ iPSCs compared to the control iPSCs. However, specific MMPs did not differ statistically (Figure 1G). ROS production was analyzed using 2’, 7’-dichlorodihydrofluorescein diacetate (DCFDA) and MitoTracker Deep Red (MTDR) double staining and flow cytometry. Alpers’ iPSCs produced more total ROS than the control iPSCs (Figure 1H). After normalizing total ROS by measuring mitochondrial mass MTDR, no difference was observed in ROS production per mitochondria (Figure 1I). Furthermore, total mitochondrial ROS measured by a mitochondrial ROS (mito-ROS)-sensitive fluorescent dye, did not differ between Alpers’ and control iPSCs (Figure 1J). When normalized by MTG, specific mito-ROS levels remained statistically indistinct (Figure. 1K).

ATP production per cell was assessed using a luminescence assay, revealing no difference in ATP levels between Alpers’ and control iPSCs (Figure 1L). Moreover, to explore glycolysis activity in Alpers’ iPSCs, we measured L-lactate levels and found them to be higher in Alpers’ iPSCs compared to the control group (Figure 1M).

To further investigate the mtDNA copy number, we employed flow cytometry to indirectly mtDNA copy number by examining the levels of mitochondrial transcription factor A (TFAM) and mtDNA, both of which are bound together in molar amounts. We also sought to determine the mtDNA copy number per mitochondria by performing double staining with the mitochondrial import receptor subunit TOM20 (TOMM20) and then calculating the ratio of TFAM to TOMM20. Our findings revealed no significant difference in the expression of TFAM and TOMM20 proteins between the patient and control (Figure. 1N-O).

Given the known association of oxidative phosphorylation (OXPHOS) complex deficiency with *POLG* mutations [30], we evaluated whether this also the case in iPSCs derived from Alpers’ iPSCs. We did this by using antibodies against the CI subunit NDUFB10 for measurement of CI level, the complex IV subunit COXIV to assess mitochondrial complex IV (CIV) level, and TOMM20 to quantify the mitochondrial volume. Following this, we applied flow cytometry.

Our findings revealed that both total levels of NDUFB10 (Figure. 1P) and specific levels of NDUFB10 when normalized to TOMM20 levels (Figure. 1Q) were significantly reduced in Alpers’ iPSCs. We did not find any significant changes in the total (Supplemental Figure 3) and specific CIV (Figure 1R) or the total TOMM20 level (Figure 1N) in Alpers’ iPSCs compared to control iPSCs.

Collectively, these results suggest that we can successfully generate iPSCs from a patient with Alpers’syndrome and that the iPSCs derived from the patient exhibit mild mitochondrial alterations, including an elevated L-lactate level and depletion of CI.

### Alpers’ NSCs showed more severe impairment of mitochondrial function compared to controls

Continuing our investigations, we utilized a method from previous research to further differentiate iPSCs into NSCs, and then conducted the same experiments as before to evaluate alterations in mitochondrial function in these NSCs. Alpers’ NSCs were observed to display a conventional neural stem cell morphology (Figure. 2A) and an unaltered mitochondrial double cristae morphology (Figure 2B). The NSCs, derived from iPSCs, showed the positive expression of neural stem cell markers, Nestin and SOX2 (Figure 2C). Utilizing flow cytometry, it was established that over 93% and 87% of Alpers’ NSCs expressed Nestin and SOX2, respectively (Figure 2D).

**Figure 2.**
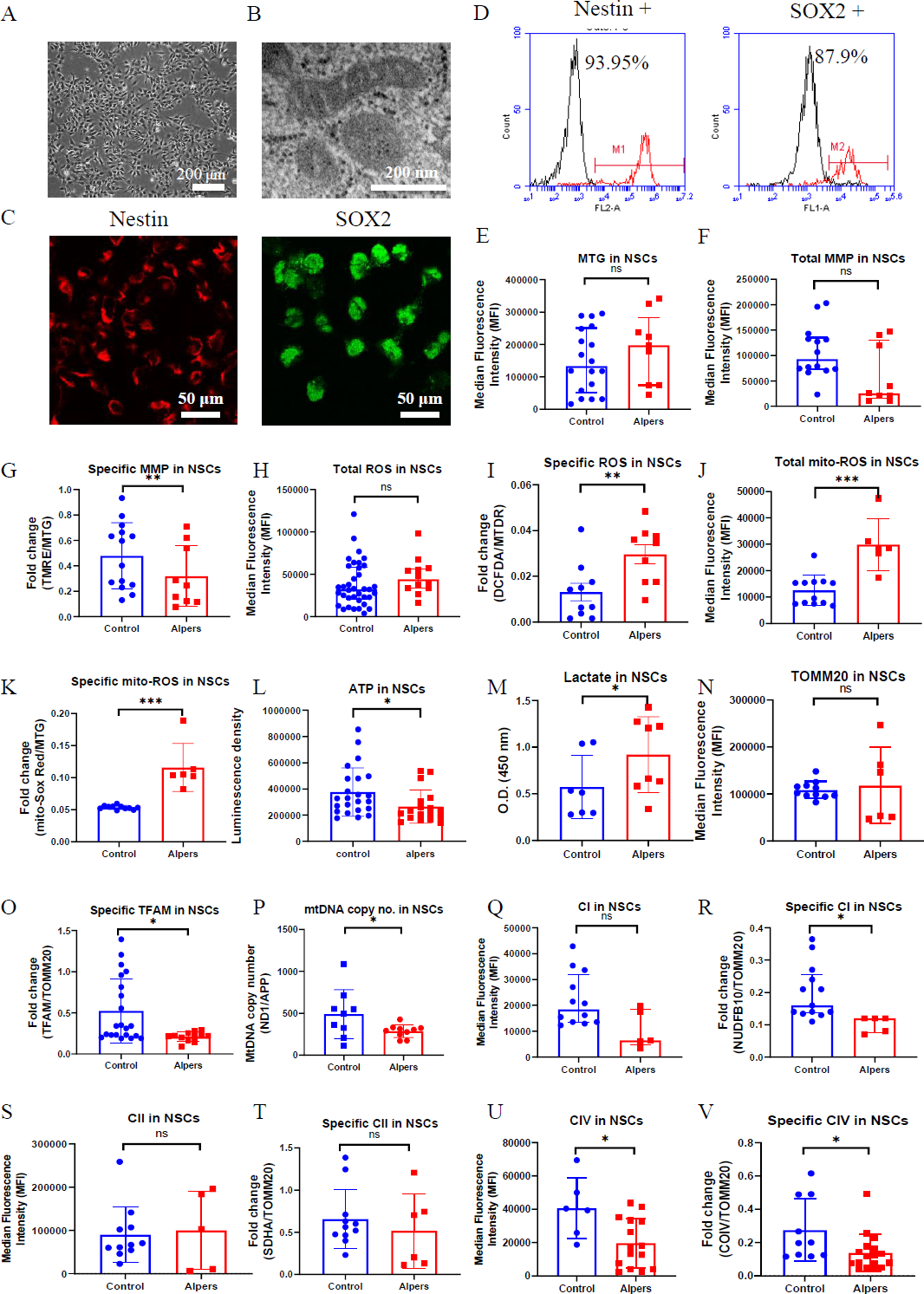
Measurement of mitochondrial function, mtDNA alteration and mitochondrial complexes in Alpers’ NSCs. A. Representative phase-contrast images of NSCs of Alpers’ patients. Scale bar is 200 µm. B. Representative TEM images of NSCs of Alpers’ patients. Scale bar is 50 µm. C. Representative fluorescent images of Alpers’ NSCs for NSC markers Nestin and SOX2. D. Flow cytometric analysis for NSC markers Nestin and SOX2 in NSCs. E. Flow cytometric analysis for mitochondrial volume using MTG in NSCs. F-G. Flow cytometric analysis for total MMP and specific MMP in NSCs. H-K. Flow cytometric analysis for total ROS, specific ROS, total mito-ROS, and specific mito-ROS in NSCs. L-M. Flow cytometric analysis for ATP and lactate production in NSCs. N. Flow cytometric analysis for TOMM20 levels in NSCs. O-P. Flow cytometric analysis for TFAM and mtDNA copy number levels in NSCs. Q-V. Flow cytometric analysis for CI, specific CI, CII, specific II, CIV, specific CIV levels in NSCs.

Using the same TMRE/MTG double staining method as we did for iPSCs, it was found that specific MMPs were decreased in Alpers’ NSCs compared to controls (Figure 2G). However, no significant variation was observed in total MMPs (Figure 2F) and mitochondrial volume, as indicated by MTG (Figure 2E). For Alpers’ NSCs, a significant increase was observed in specific intracellular ROS (DCFDA/MTDR) and mitochondrial ROS at both total (MitoSOX Red) and specific level (MitoSOX Red/MTG) when compared to control NSCs (Figure. 2I-K), yet the total intracellular ROS (DCFDA/MTDR) showed no difference (Figure 2H). A direct measurement of intracellular ATP production revealed a marked decrease in ATP production in Alpers’ NSCs compared to the control NSCs (Figure 2L), whereas the L-lactate levels were found to be higher in Alpers’ NSCs than in controls (Figure 2M).

In addition, we utilized indirect methods TFAM measurement and qPCR to evaluate the mtDNA levels of ND1/APP. Using flow cytometry, we discerned a marked decline in specific TFAM levels (TFAM/TOMM20) in Alpers’ NSCs compared to controls (Figure 2N). Interestingly, we didn’t observe any significant alterations in the levels of TOMM20 (Figure 2O). To corroborate the reduction in the mtDNA levels as observed through the ND1/APP measurements in Alpers’ NSCs, we performed qPCR (Figure 2P), which confirmed the same. Further assessments of OXPHOS complexes I, II, and IV using flow cytometry revealed significantly decreased expression, both in terms of the total level and specificity of CI (Figure 2Q, R) and CIV (Figure 2S, T) in Alpers’ NSCs compared to control NSCs. However, we did not notice any significant changes in the total (Figure 2U) and specific level of complex II (CII) (Figure 2V).

This body of evidence collectively suggests that NSCs in Alpers’ syndrome are distinguished by profound mtDNA depletion and resultant mitochondrial dysfunction.

### Mitochondrial-related pathways were downregulated in Alpers’ iPSCs and NSCs, and more pronounced in NSCs

In order to delve deeper into the comparison of transcriptome and gene alterations between Alpers’ and control lines, we executed a transcriptome-wide RNA-seq analysis on both iPSCs and induced NSCs derived from Alpers’ and controls. Hierarchical clustering based on RNA transcripts displayed unique, independent clusters for NSCs and iPSCs (Figure 3A). Principal component analysis (PCA) authenticated the separation between NSCs and iPSCs, with the difference between Alpers’ NSCs and control NSCs being more conspicuous than those between iPSCs (Figure 3B). The differential expression analysis unveiled that 1805 genes were upregulated, and 1905 genes were downregulated in Alpers’ iPSCs compared with the control iPSCs (Figure 3C).

**Figure 3.**
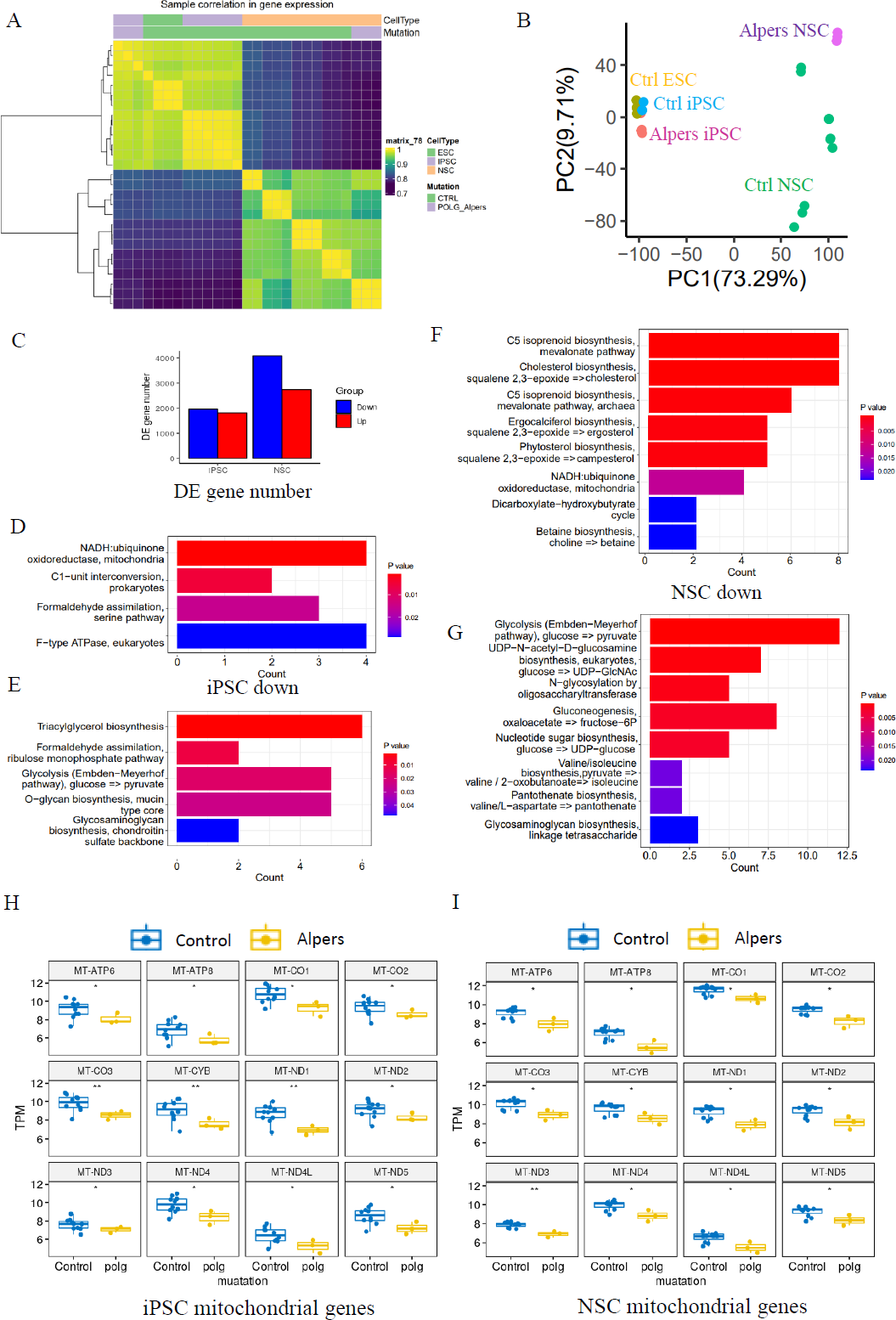
Transcriptomic status and dysregulated mitochondrial related pathways of Alpers’ patients derived iPSCs and NSCs. A. Sample correlation in transcriptomic status of Alpers’ patient derived iPSCs and NSCs, and the corresponding controls. B. PCA analysis of transcriptomic expression data for Alpers’ patient derived iPSC and NSCs and controls. C. Number of upregulated and down-regulated DE genes between Alpers’ and NSCs in iPSCs and NSCs, respectively. D. Downregulated metabolic pathways in Alpers’ iPSCs compared with controls. E. Upregulated metabolic pathways in Alpers’ iPSCs compared with controls. F. Upregulated metabolic pathways in Alpers’ NSCs compared with controls. G. Downregulated metabolic pathways in Alpers’ iPSCs compared with controls. H-I. RNA expressions of mitochondrial genes in Alpers’ iPSCs and NSCs.

Relative to controls, Alpers’ NSCs displayed 2725 upregulated genes and 4059 downregulated genes (Figure 3C). KEGG pathway analysis based on DEGs disclosed a downregulation of the mitochondrial NADH: ubiquinone oxidoreductase pathway in both Alpers’ derived iPSCs (Figure 3D, E, and Table S1) and NSCs (Figure 3F and Table S2) compared to control cells. Interestingly, gluconeogenesis and glycolysis pathways were markedly upregulated in Alpers’ NSCs (Figure 3G and Table S2), indicating that glycolysis might be compensating for energy production in Alpers’ NSCs, a phenomenon providing indirect evidence of mitochondrial dysfunction. Moreover, Alpers’ iPSCs and NSCs exhibited significantly lower levels of mitochondrial transcripts, encompassing *ATP6*, *ATP8*, *CO1*, *CO2*, *CO3*, *CYB*, *ND1*, *ND2*, *ND3*, *ND4*, *ND4L*, and *ND5*, relative to controls (Figure 3H, I). This observation aligns with the previously demonstrated mtDNA depletion in NSCs derived from Alpers’ patients (Figure 2O, P).

Taken together, this body of evidence proposes that mitochondrial-associated pathways are downregulated in both iPSCs and NSCs derived from Alpers’ patients, with these alterations being more accentuated in NSCs.

### Alpers’ cortical organoids demonstrated cortical neuronal loss and astrocyte accumulation

In our next step, we delved deeper into the mitochondrial changes in 3D cortical brain development, employing a cortical organoid model. This is a relatively complex 3D model that more accurately replicates the microenvironment and structure of the human brain (Figure 4A). Using a previously documented procedure, we transformed iPSCs into cortical organoids (Figure 4B) [31]. By days 40-50, these cortical organoids had matured into large (2-3 mm in diameter) and complex heterogeneous tissues (Figure 4C). At day 40, brain organoids presented intricate morphology, including cortical folded surfaces and sulci, and expressed the mature neural marker MAP2 (Figure 4D), neonatal neural marker tubulin β III (Tuj1), ventricular zone (VZ) marker SOX2 (Figure 4E and Supplemental S4). By day 90, a majority of neurons in the organoids were positive for mature neural marker NeuN in outer layers (Figure 4F and Supplemental Figure 5). Additionally, a minor fraction of cells in the organoids were GFAP positive (Figure 4G, F and Supplemental Figure 5). Furthermore, the cortical pyramidal neuronal marker CTIP2 and SATB2 were also expressed in the upper layers and middle layer of the organoids (Figure 4G and Supplemental Figure 5), demonstrating that the organoids’ resemblance to cortical neurons.

**Figure 4.**
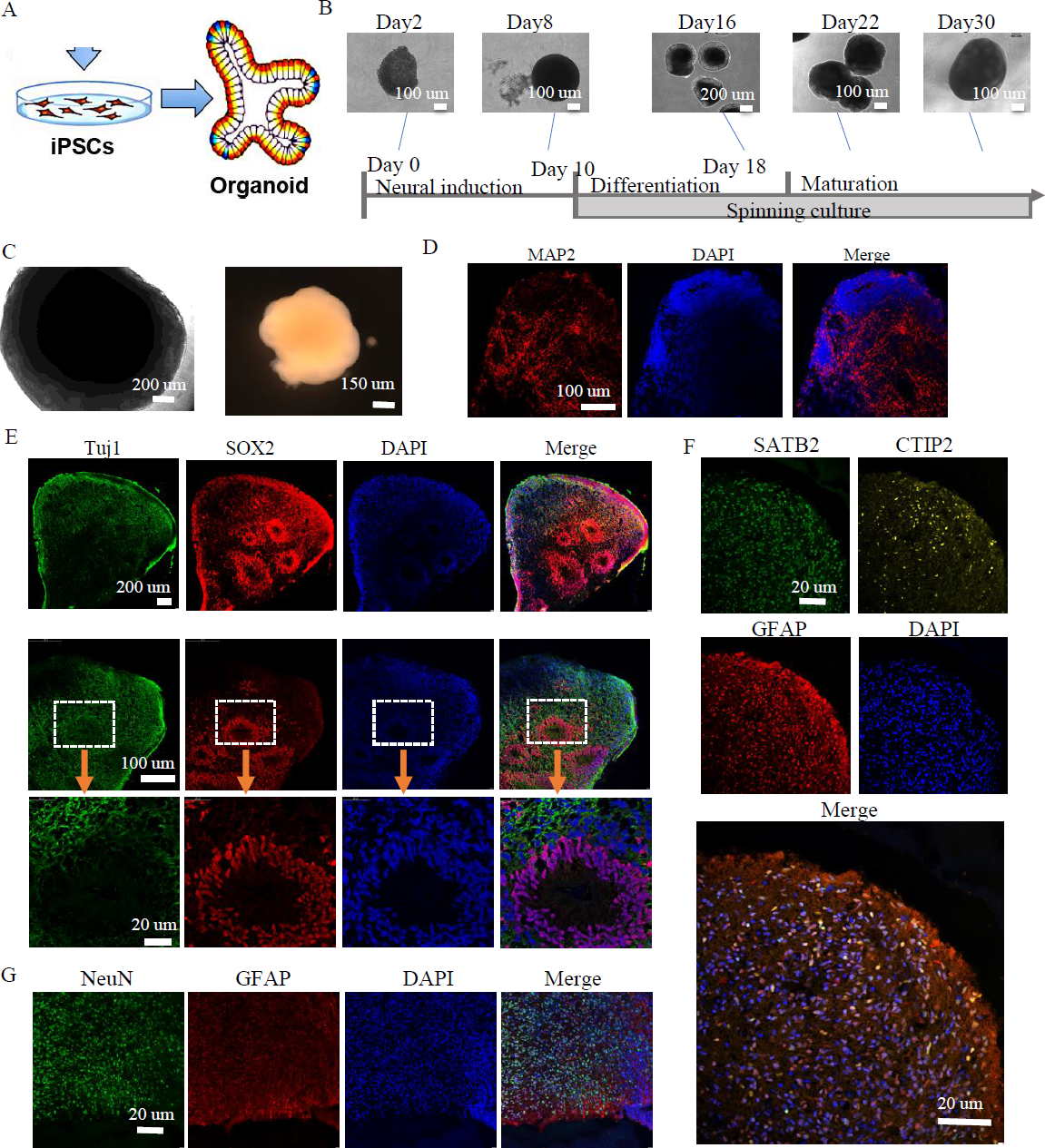
Generation of cortical brain organoid from iPSCs. A. Illustration of generation of cortical brain organoid. B. Generation of cortical brain organoid using iPSCs. Scale bar is 100 µm or 200 µm. C. Phase-contrast images of cortical organoids of Alpers’ patient and controls at day 54. Scale bar is 200 µm or 150 µm. D. Fluorescent staining of cortical organoid section using mature neuron marker MAP2 at day 40. Nuclei are stained with DAPI (blue). Scale bar is 20 µm. E. Fluorescent staining of cortical organoid section using neuron marker Tuj1 and neural progenitor marker SOX2 at day 40. Nuclei are stained with DAPI (blue). Scale bar is 20 µm. F. Fluorescent staining of cortical organoid section using neuron marker NeuN and astrocyte marker GFAP at day 90. Nuclei are stained with DAPI (blue). Scale bar is 20 µm. G. Fluorescent staining of cortical organoid section using cortical neuronal markers SATB2, CTIP2 and astrocyte marker GFAP at day 90. Nuclei are stained with DAPI (blue). Scale bar is 20 µm.

We then generated cortical organoids from iPSCs from Alpers’ patients. Compared to controls, these organoids exhibited irregular morphology and folding patterns (Figure 5A and Supplemental Figure 6). Upon performing immunofluorescent staining of 70-day-old cortical organoids, we found that while normal control cortical organoids formed a typical cortical layer structure characterized by prominent cortical neurons, including SATB2- and CTIP2-positive neurons, Alpers’ organoids exhibited irregular folding without typical cortical layers (Figure 5A and Supplemental Figure 6). This suggested abnormal cortical development in Alpers’ organoids (Figure 5B). Further quantification of SATB2, CTIP2 cortical neurons, and MAP2-positive mature neurons in mature cortical organoids (day 90) revealed significantly lower expression of SATB2 and CTIP2 in Alpers’ organoids (Figure 5C, G-H). The level of MAP2 expression was also lower in Alpers’ organoids than in controls (Figure 5C, F), confirming a loss of cortical neurons in Alpers’ cortical organoids. Meanwhile, we observed an increase in SOX2-positive neural progenitors in Alpers’ organoids compared with controls (Figure 5D, K). Previous studies have indicated that loss of mtDNA leads to a reactive increase in astrocytes [32]. To further explore an astrocyte differentiation in Alpers’ organoids, we examined the number of astrocytes in mature cortical organoids (day 90) and found that the number of GFAP-positive astrocytes in Alpers’ organoids was significantly higher than in controls (Figure 5D, J). This suggests that astrogliosis accompanying h cortical neuronal loss is a characteristic of cortical brain pathology in Alpers’ disease. Additionally, we used immunofluorescence to measure the CI subunit NDUFB10 and discovered significantly reduced levels in Alpers’ organoids compared to the control (Figure 5E, L-M). These results affirm our ability to differentiate 3D cortical organoids from iPSCs derived from Alpers’ patients, demonstrating that these organoids recapitulate disease-specific pathological and molecular features similar to those observed in the patient’s brain.

**Figure 5.**
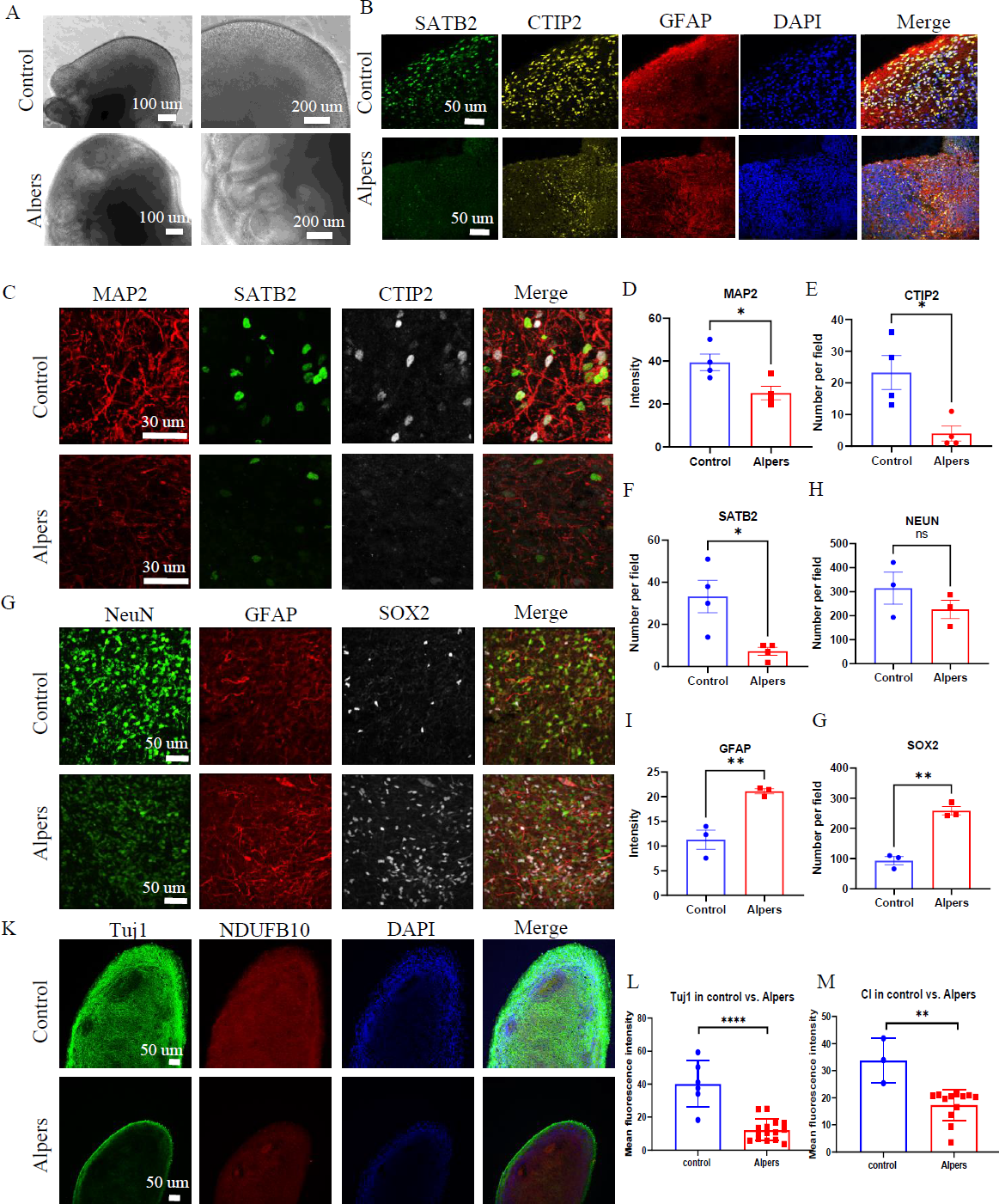
Cortical layer deformation, neuronal loss and astrocyte proliferation in Alpers’ cortical organoid. A. Phase-contrast images of cortical organoids of Alpers’ patient and controls at day 90. Scale bar is 100 µm or 200 µm. B. Fluorescent staining of Alpers’ and control cortical organoid section using cortical neuronal markers SATB2, CTIP2 and astrocytes marker GFAP at day 90. Nuclei are stained with DAPI (blue). Scale bar is 50 µm. C. Fluorescent staining of Alpers’ and control cortical organoid (day 90) section using neural fiber marker MAP2 and cortical neuronal markers SATB2, CTIP2. Nuclei are stained with DAPI (blue). Scale bar is 30 µm. D. Fluorescent staining of Alpers’ and control cortical organoid (day 90) section using astrocyte marker GFAP, neural marker NEUN, and neural progenitor marker SOX2. Nuclei are stained with DAPI (blue). Scale bar is 50 µm. E. Fluorescent staining of Alpers’ and control cortical organoid (day 90) section using mitochondrial CI marker NDUFB10 and neural marker Tuj1. Nuclei are stained with DAPI (blue). Scale bar is 50 µm. F-M. Quantification of immunofluorescent staining of MAP2, CTIP2, SATB2, NEUN, GFAP, SOX2, Tuj1 and CI in Alpers’ cortical organoids compared with controls. Nuclei are stained with DAPI (blue). Scale bar is 50 µm.

### Alpers’ cortical organoids exhibited significant downregulation of mitochondrial and synaptogenesis-related pathways, while there was an upregulation of pathways associated with astrocyte/glial cells and neuroinflammation

To further explore regulatory pathways in Alpers’ cortical organoids, we performed RNA sequencing analysis on both Alpers’ and control organoids. We selected 90-day-old cortical organoids for this analysis because, as mentioned earlier, they display a relatively mature state with a layered structure of cortical neurons and glial cells (Figure 4F, G and Supplemental Figure 4). The PCA analysis demonstrated that Alpers’ and control organoids formed distinct clusters (Figure 6A), suggesting a divergence in expression profiles between the two groups. DEGs analysis revealed 1440 differentially expressed genes, with 1010 genes being up-regulated and 430 genes down-regulated in Alpers’ organoids compared to the control group (Figure 6B). Furthermore, when comparing the expression of mitochondrial transcripts in the Alpers’ group to the control group, significant decreases in the expressions of *MT-CO2*, *MT-CYB, MT-ND1, MT-ND4, MT-ND4L,* and *MT-ND5* were noted in Alpers’ organoids (Figure 6C).

**Figure 6.**
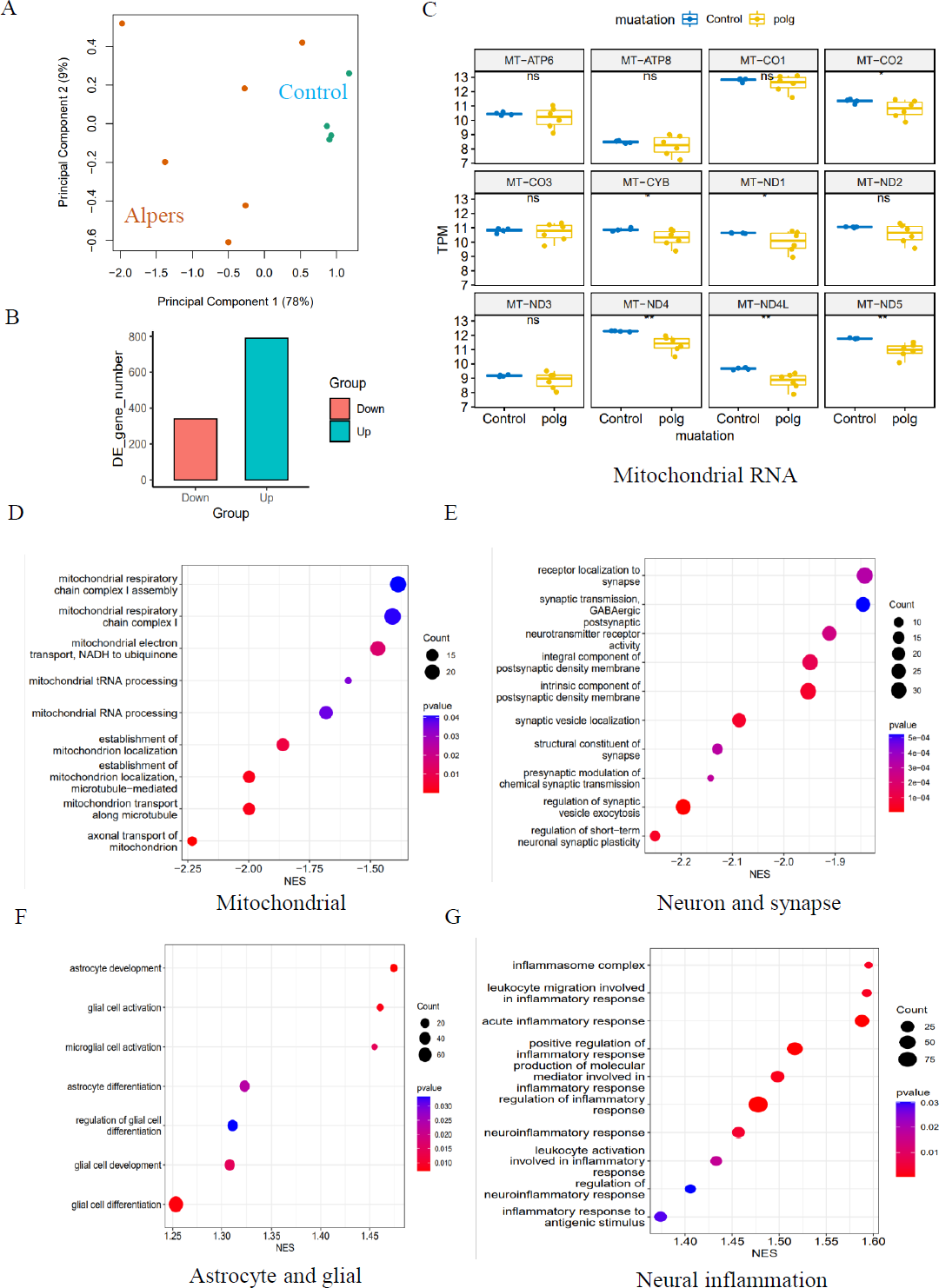
Transcriptomic status and dysregulated pathways of Alpers’ patients derived cortical organoids. A. PCA analysis of transcriptomic expression data for Alpers’ patient derived cortical organoids and controls. B. Number of upregulated and down-regulated DE genes in Alpers’ cortical organoids compared with controls. C. RNA expressions of Mitochondrial encoded genes in Alpers’ cortical organoids. D. Mitochondrial related pathways were downregulated in Alpers’ cortical organoids. E. Synapse maturation related pathways were downregulated in Alpers’ cortical organoids. F. Astrocytes and glial related pathways were upregulated in Alpers’ cortical organoids. G. Neural inflammation related pathways were upregulated in Alpers’ cortical organoids.

Gene set enrichment analysis (GSEA) results indicated that Alpers’ organoids exhibited substantial downregulation of mitochondrial and synaptogenesis-related pathways. This includes mitochondrial-related pathways including mitochondrial CI, mitochondrial electron transport, mitochondrial tRNA processing, and mitochondrial localization and trafficking (Figure 6D and Table S3), as well as axonal trafficking and synapse-related pathways including presynaptic transmission, synaptic vesicle exocytosis, and postsynaptic receptor activity (Figure 6E and Table S4). GABAergic neurons, inhibitory interneurons crucial in preventing epilepsy [33], showed marked downregulation of their synaptic transmission pathway in Alpers’ cortical organoids (Figure 6E and Table S4), indicating pathological changes of GABAergic neurons in Alpers’ disease. Moreover, astrocyte- and glial-related pathways were upregulated in Alpers’ cortical organoids (Figure 6F and Table S5), consistent with the immunostaining findings described earlier (Figure 5D, J). Intriguingly, cortical organoids from Alpers’ patients also demonstrated an up-regulation of neuroinflammatory pathways (Figure 6G and Table S6).

These findings suggest that Alpers’ cortical organoids display a distinct transcriptomic profile, characterized by the downregulation of mitochondrial and synaptogenesis-related pathways and upregulation of astrocyte/glial-related and neuroinflammatory pathways.

### Long-term treatment with NR partially ameliorated the neurodegenerative alterations observed in Alpers’ cortical organoids

NAD^+^ is a crucial metabolite that plays pivotal roles in cellular energy metabolism, genomic stability, and mitochondrial balance, with impacts on neurogenesis and the organization of neuronal networks [34]. Prior research has highlighted that the enhancement of NAD^+^ levels through supplementation can significantly bolster mitochondrial function, positioning NAD^+^ as a promising neuroprotective agent [35]. In alignment with these findings, our results illustrated a severe disruption of the NADH pathway and mitochondrial function in the iPSC-NSCs model of Alpers’ syndrome compared to the control group.

To test the potential of NAD^+^ supplementation in mitigating this dysfunction, we administered NR, a form of NAD^+^, to Alpers patient-derived cortical organoids that were one month old, daily over a two-month period (Figure 7B). We observed that the NR-treated organoids exhibited increased expression of MAP2 compared to those untreated (Figure 7C, F), alongside an elevation in the cortical neural markers SATB2 and CTIP2 (Figure 7C, G-H). Interestingly, a decrease in the number of GFAP-positive astrocytes was noticed post NR treatment when compared to untreated organoids (Figure 7D, J). Immunostaining of Complex I subunit NDUFB10 showed a marked rise in NDUFB10 expression after the treatment with NR (Figure 7K, M).

**Figure 7.**
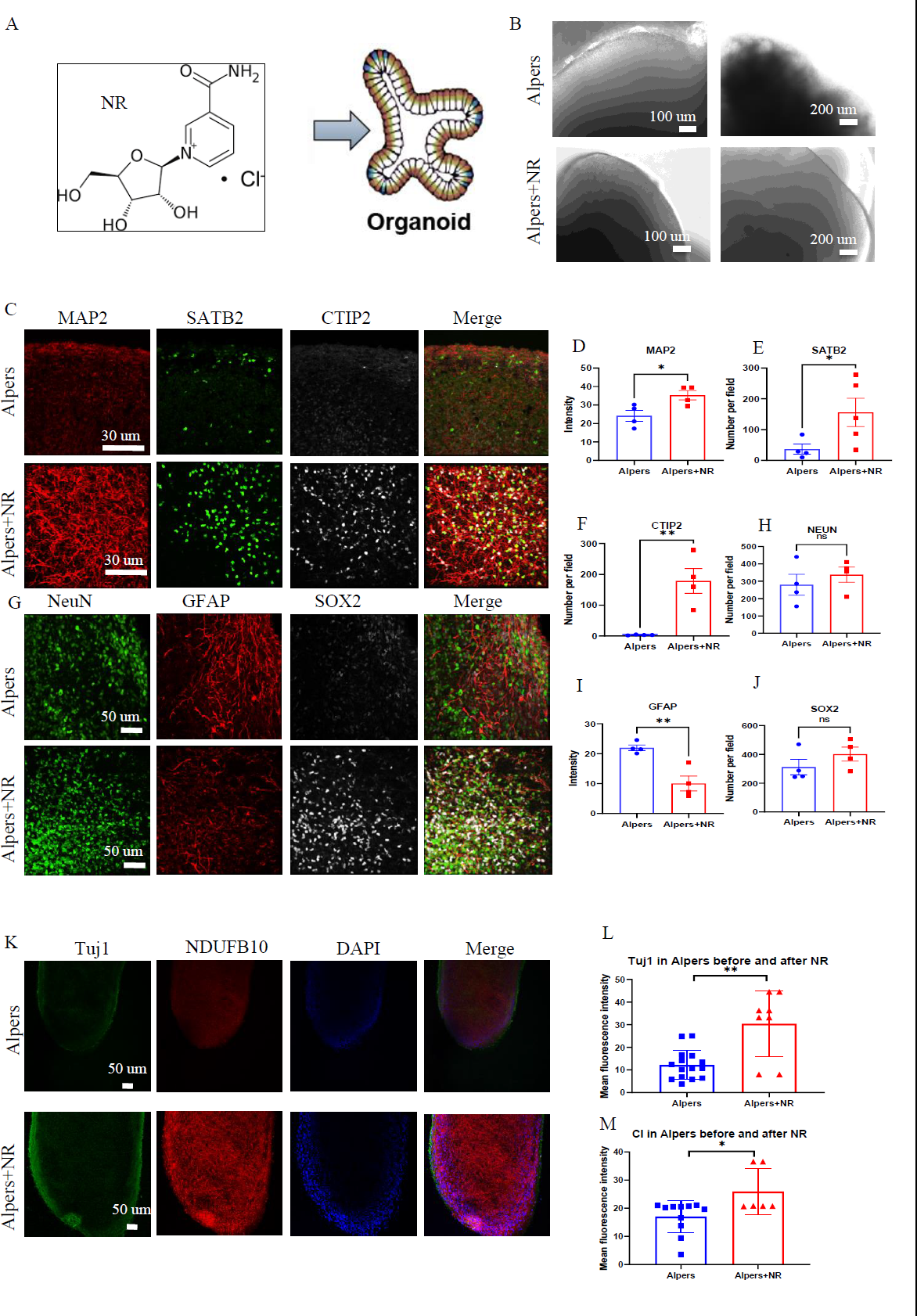
Long time treatment of NR partially rescues changes of Alpers’ cortical organoids. A. Illustration of treatment of NR of Alpers’ cortical organoids. B. Phase-contrast images of NR treatment of cortical organoids of Alpers’ patients. Scale bar is 100 µm or 200 µm. C. Fluorescent staining of NR treated Alpers’ cortical organoid section using neural fiber marker MAP2 and cortical neuronal markers SATB2, CTIP2. Nuclei are stained with DAPI (blue). Scale bar is 30 µm. D-F. Quantification of immunofluorescent staining of MAP2, SATB2 and CTIP2 in NR treated Alpers’ cortical organoids compared with organoids without treatment. G. Fluorescent staining of NR treated Alpers’ cortical organoid section using astrocyte marker GFAP, neural marker NeuN, and neural progenitor marker SOX2. Nuclei are stained with DAPI (blue). Scale bar is 50 µm. H-J. Quantification of immunofluorescent staining of NeuN, GFAP and SOX2 in NR treated Alpers’ cortical organoids compared with organoids without treatment. K. Fluorescent staining of NR treated Alpers’ cortical organoid section using complex I marker NDUFB10 and neural marker Tuj1. Nuclei are stained with DAPI (blue). Scale bar is 50 µm. L-M. Quantification of immunofluorescent staining of Tuj1 and complex I marker NDUFB10 in NR treated Alpers’ cortical organoids compared with organoids without treatment.

An RNA sequencing analysis was performed to identify genetic changes before and after NR treatment. PCA demonstrated that NR treatment altered the RNA expression profile of Alpers’ organoids significantly, moving it closer to that of healthy controls (Figure 8A). GSEA showed that NR treatment was able to reverse the dysregulated pathways that were identified in Alpers’ patient organoids. While mtDNA RNA levels did not see a significant rise with NR treatment (Supplemental Figure 7), we noted an up-regulation in the activity of pathways related to mitochondrial function, such as mitochondrial electron transport, proton transport ATP synthase complex, mitochondrial transport along microtubules and axons, and mitophagy in response to mitochondrial depolarization (Figure 8B and Table S7).

**Figure 8.**
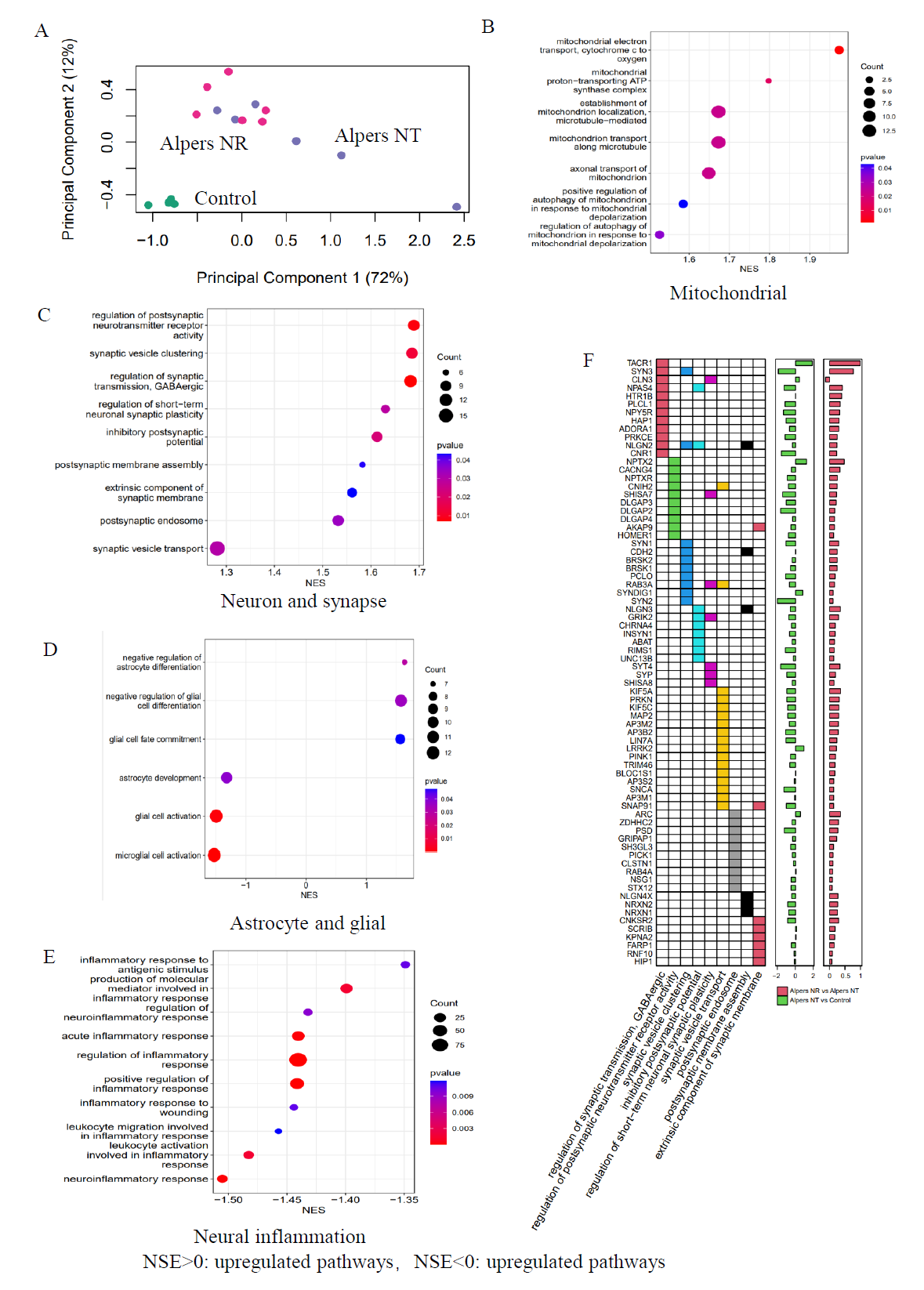
Transcriptomic data shows the treatment effect of NR on Alpers’ cortical organoids. A. PCA analysis of transcriptomic expression data for NR, non-NR treated Alpers’ cortical organoid and healthy controls. B. Mitochondrial related pathways were upregulated in NR treated Alpers’ cortical organoids. C. Synapse maturation related pathways were upregulated in NR treated Alpers’ cortical organoids. D. Astrocytes and glial related pathways were downregulated in NR treated Alpers’ cortical organoids. E. Neural inflammation related pathways were downregulated in NR treated Alpers’ cortical organoids. F. Heatmap showing the genes that were involved in significantly enriched synapse formation related pathways in NR treated Alpers’ cortical organoids.

Further, patient organoids treated with NR demonstrated an upregulation of pathways associated with synapses, including postsynaptic pathways (membrane assembly, endosomal and neurotransmitter receptor activity), inhibitory neuronal pathways (regulator of GABAergic synaptic transmission, inhibitory postsynaptic potentials), and synaptic vesicles (synaptic vesicle aggregation and trafficking) (Figure 8C, F and Table S8). In contrast, astrocyte and glial cell pathways were downregulated after NR treatment, including astrocyte development, glial cell activation, glial cell fate commitment, and pathways negatively regulating astrocyte/glial cell differentiation (Figure 8D and Table S9). Intriguingly, we also observed a downregulation of pathways related to neuroinflammation following NR treatment (Figure 8E and Table S10).

Taken together, these findings suggest that the supplementation of NAD^+^, specifically in the form of NR, could be an effective therapeutic strategy to counteract neuronal loss, glial enrichment, and mitochondrial damage observed in the cortex of Alpers’ syndrome patients.

## Discussion

Alpers’ syndrome, the most severe condition related to *POLG* mutations, primarily impacts the nervous system and liver. This syndrome stems from a genetic anomaly that triggers an enzymatic change, leading to mtDNA reduction and the onset of mtDNA depletion syndrome. Our understanding of this disease and the development of effective treatments have been hindered by the limited access to human tissue and the absence of reliable disease models. However, the ability to reprogram patient cells and differentiate them into the affected cell types has been transformative for the field [36, 37]. In this study, we reprogrammed Alpers’ patient fibroblasts carrying compound heterozygous *POLG* mutations A467T (c.1399G>A), P589L (c.1766C>T) into iPSCs, and subsequently differentiated them into NSCs and 3D cortical organoids These patient specific NSCs mimicked the mtDNA depletion and CI deficiency seen in postmortem tissue. Similarly, the 3D cortical organoids manifested changes akin to those observed in the patient’s cerebral cortex, including cortical neuronal loss and astrogliosis [38]. This work represents the first instance of accurately replicating the pathological changes seen in autopsy studies using an iPSC-derived cell model from Alpers’ patients. Notably, by employing iPSC derived NSCs, we managed to pinpoint cellular and molecular mechanisms involved in the disease process. These mechanisms included the overproduction of ROS linked to suppressed CI-driven respiration and defective NAD^+^ metabolism. Additionally, by supplementing NAD^+^, we managed to alleviate the changes in cortical areas. Our findings not only offer compelling evidence of the various cellular and molecular mechanisms accountable for the reduced neuronal pools in POLG-related diseases, but also pave the way for the development of novel diagnostic tools and therapeutic interventions.

We found that Alpers’ iPSCs exhibited reduced CI levels, while maintaining MMP, mitochondrial ROS and ATP production. Coupled with our finding of decreased mtDNA transcript levels in Alpers’ iPSCs, this data suggests that mtDNA depletion in these Alpers’ does not result in significant mitochondrial dysfunction. One possible explanation for this observation is iPSCs’ higher reliance on glycolysis for iPSC energy productivity than oxidative phosphorylation [39].

Given that neuronal cells, including NSCs, rely more heavily on mitochondrial OXPHOS for their energy needs [40], iPSCs from Alpers’ manifested a more pronounced impairment of mitochondrial function upon differentiated into neurons. In these Alpers’ NSCs we observed a lower MMP, decreased ATP production, and elevated ROS generation. We also noted significant reductions in mitochondrial CI, III, IV protein levels and mtDNA transcripts. Our research thus illustrates that the impacts of mtDNA loss in Alpers’ mutations start to become evident upon differentiation into NSCs. This indicates that NSCs serve as an effective model for studying diseases related to Alpers’ mutations. As NSCs have the capacity to differentiate into a wide array of neuronal and glial cell lineages, mitochondrial dysfunction in NSCs might carry implications for the differentiation and development of various neuronal and glial cell lines.

The key clinical manifestations of Alpers’ syndrome typically include refractory epilepsy, cognitive decline, and developmental regression, predominantly affecting children 6 months to 3 years of age. Pathological investigations have revealed that neuronal degeneration and cortical laminar necrosis serve as the primary contributors of neuronal loss and cerebral atrophy in Alpers’ patients [41]. Upon our examination of substantial mitochondrial dysfunction in patient derived NSCs, we proceeded to generate cortical organoids from Alpers’ patients. Our observations revealed that these cortical organoids primarily exhibited a disarray in cortical layering along with a notable loss of SATB2 and CTIP2 cortical neurons. The analysis of transcriptomic data disclosed a significant decrease in the expression of mtDNA transcripts within Alpers’ organoids and the downregulation of mitochondrial-associated pathways, findings that align with those derived from NSCs. Of particular note, neuron- and synapse-related pathways demonstrated significant down-regulation, with GABAergic inhibition of neuron-related synaptic pathways being particularly affected. This suggests that neuronal interconnections may be compromised in Alpers’ syndrome, a conclusion that aligns with previous findings regarding brain pathology in Alpers’ patients [41]. These results propose that cortical neuronal loss, as a secondary effect to mtDNA depletion and compromised ATP export in Alpers’ patients, might contribute to central nervous system symptoms observed in these patients, including epilepsy and dementia. A potential extension of our research could include an exploration of the specific molecular mechanisms contributing to the downregulation of neuron- and synapse-related pathways in Alpers’ patients. Future studies could also delve into the potential for therapeutic interventions that might bolster these pathways, potentially mitigating some of the neurological impacts of the disease. Furthermore, it may be insightful to examine whether these impacts are consistent across different ages, genders, and disease progression stages, or if distinct trends emerge.

The irregular morphology, atypical cortical layers, and irregular folding of Alpers’ organoids underscore aberrant cortical development, potentially contributing to the clinical manifestation of Alpers’ syndrome. Further substantiating this, the organoids showed a significant reduction in the expression of cortical neuronal markers, SATB2 and CTIP2, and mature neural marker, MAP2, signifying the loss of cortical neurons. Additionally, the overexpression of SOX2-positive neural progenitors and an increased number of GFAP-positive astrocytes point towards astrogliosis in the context of cortical neuronal loss, a hallmark of brain pathology in Alpers’ syndrome .These findings align with the understanding that Alpers’ syndrome is a neurodegenerative disorder characterized by the premature loss of neurons, particularly within the cortex, and the concurrent proliferation of astrocytes (a process known as astrogliosis) in response to neuronal damage [42]. Further, the reduced expression of CI subunit NDUFB10 strengthens the notion of impaired mitochondrial function in Alpers’ disease, which aligns with findings from previous studies conducted on postmortem brain tissue [41].

The differential gene expression observed through RNA sequencing analysis gives a more detailed view of the altered molecular landscape within Alpers’ organoids. PCA revealed a clear distinction in gene expression profiles between Alpers’ and control organoids. This finding is consistent with previous research highlighting the distinct genetic signatures associated with neurodegenerative disorders [43], including Alpers’ syndrome. Of particular note, the downregulation of mitochondrial transcripts *MT-CO2*, *MT-CYB*, *MT-ND1*, *MT-ND4*, *MT-ND4L*, and *MT-ND5* implies impaired mitochondrial function in Alpers’ disease, in agreement with the postulated role of mitochondrial dysfunction in this condition [41]. The GSEA further uncovered dysregulated pathways in Alpers’ organoids. The downregulation of mitochondrial and synaptogenesis-related pathways, including presynaptic transmission, synaptic vesicle exocytosis, and postsynaptic receptor activity, could underlie the neurological symptoms seen in Alpers’ syndrome, such as seizures, developmental regression, and cognitive decline. This is especially evident with the observed downregulation of GABAergic synaptic transmission pathway, given the key role of GABAergic interneurons in epilepsy pathogenesis [44]. In contrast, the upregulation of astrocyte- and glial-related pathways, coupled with the evidence of increased astrocyte differentiation, suggests reactive astrogliosis in Alpers’ syndrome. Furthermore, the upregulation of neuroinflammatory pathways may reflect an innate immune response to ongoing neuronal damage and loss, an aspect that warrants further exploration. Taken together, the insights derived from these 3D cortical organoids offer a promising avenue for exploring the complex pathophysiology of Alpers’ syndrome, facilitating the development of potential therapeutic interventions. Moreover, the upregulation of neuroinflammatory pathways could represent a novel target for therapeutic strategies, given the increasing evidence for the involvement of neuroinflammation in neurodegenerative disorders [45, 46]. However, more work is needed to translate these findings into practical applications. In addition to studying these mechanisms in more detail, future research might focus on testing potential treatments in this model or finding ways to correct the observed dysregulations, possibly through gene therapy or other innovative techniques.

In this study, the role of NAD^+^ supplementation, specifically the administration of NR, was investigated as a potential therapeutic strategy to alleviate the pathological changes in Alpers’ syndrome. NAD^+^ is a crucial metabolite with a myriad of roles within the cell, including functions in cellular bioenergetics, genome stability, and mitochondrial homeostasis, all processes that are perturbed in Alpers’ syndrome. NR supplementation has previously been shown to improve mitochondrial function [47], restore NAD^+^ levels, and have a neuroprotective effect [48]. These findings provided the rationale for investigating the potential benefits of NR supplementation in an Alpers’ patient-derived iPSC model, which revealed profound mitochondrial dysfunction. Upon administering NR to the patient-derived cortical organoids, notable improvements were observed. Specifically, an increased expression of mature neuronal markers (MAP2) and cortical neural markers (SATB2 and CTIP2) was seen, indicating a recovery of neuronal populations in the NR-treated organoids. Additionally, a decrease in the number of GFAP-positive astrocytes, a marker for astrogliosis, was observed, further substantiating the neuroprotective effects of NR [45]. These morphological changes corresponded with the recovery of mitochondrial function, as evidenced by the increased expression of the CI subunit NDUFB10 in NR-treated organoids.

An RNA sequencing analysis offered further insights into the molecular underpinnings of NR’s therapeutic effect. PCA analysis revealed that the entire RNA expression profile of the NR-treated Alpers’ organoids shifted closer to that of the healthy controls, implying that NR treatment significantly altered gene expression. The GSEA pathway analysis corroborated these results, demonstrating upregulation of mitochondrial-related pathways and synaptic pathways, and downregulation of astrocyte/glial cell pathways and neuroinflammatory pathways in the NR-treated organoids. These findings lend credence to the potential therapeutic efficacy of NAD^+^ supplementation in Alpers’ syndrome. Collectively, these data suggest that NAD^+^ supplementation, through NR treatment, can ameliorate the cortical changes characteristic of Alpers’ syndrome, such as neuronal loss, glial enrichment, and mitochondrial damage. However, it is crucial to validate these findings through further preclinical and clinical studies. Additionally, it is necessary to establish the safety and optimal dosing of NR supplementation, considering the complexity and severity of Alpers’ syndrome.

In discussing the limitations of this study, it’s worth emphasizing the challenges posed by the inherent rarity of Alpers’ syndrome. Acquiring a large sample size of patient-derived iPSC lines for our experiments was a significant hurdle due to the scarcity of patients, coupled with the complexities involved in reprogramming patient fibroblasts into iPSCs. Out of the two patient iPSC lines initially available to us, one demonstrated poor growth during its differentiation into brain organoids, likely an outcome of impaired mitochondrial function. This line failed to form embryoid bodies during the early stages of organoid differentiation and was unable to withstand the cryopreservation process. Consequently, our experiments were conducted on two individual clones.

Furthermore, we understand the critique raised concerning the inclusion of additional patient lines or isogenic controls to bolster our conclusions. To this end, we attempted to use CRISPR on the patient’s iPSCs to generate isogenic controls. Unfortunately, the patient’s iPSCs exhibited suboptimal growth conducive to this procedure. As a solution, we decided to incorporate two different controls with multiple clones for each line, culminating in a total of four clones for the analysis. While this somewhat addressed the issue, it is nevertheless a limitation of our study, and results should be interpreted with this in mind.

Thus, although our study has yielded significant insights into the pathophysiological progression of Alpers’ disease and potential therapeutic pathways, it is important to consider the constraints tied to the limited number of patient iPSC lines and the absence of isogenic controls. Future research may benefit from advancements in methodologies and techniques that allow for more efficient generation and growth of patient-derived iPSCs, as well as more successful incorporation of isogenic controls. Despite these limitations, we believe our study offers a meaningful contribution to the understanding and potential treatment of Alpers’ disease.

In conclusion, our research underscores the applicability of iPSC derived NSC and cortical organoid models as representative systems to simulate the pathophysiological progression of Alpers’ disease. These platforms stand as robust tools for facilitating the discovery of therapeutic strategies for such debilitating conditions. Moreover, we provide evidence that NAD^+^ supplementation, specifically through NR therapy, could present a promising clinical avenue for neuroprotection not only in Alpers’ disease, but also in a broader spectrum of mitochondrial disorders.

## Materials and Methods

### Ethics Approval

The project was approved by the Western Norway Committee for Ethics in Health Research (REK nr. 2012/919).

### Generation of iPSCs and NSCs

The fibroblasts from one Alpers’ patient carrying compound heterozygous POLG mutations A467T (c.1399G>A), P589L (c.1766C>T) were collected by punch biopsy. Detroit 551 fibroblasts (ATCC CCL 110TM) and CRL2097 fibroblasts (ATCC CRL-2097™) were used as control individual. To generate iPSCs, skin fibroblasts were infected with Sendai virus vectors containing coding sequences of human OCT4, SOX2, KLF4, and c-MYC as described previously [30, 49]. All iPSC lines were maintained under feeder-free conditions using Geltrex (Life Technologies) in E8 medium (Life Technologies) in 6-well plates (Thermo Fisher Scientific). Each line was passaged with 0.5 mM EDTA (Life Technologies) at 70-80% confluency. The E8 was changed every day, and the cells passaged every 3-4 days. All the cells were monitored for mycoplasma contamination regularly using MycoAlert™ Mycoplasma Detection Kit (Lonza). The process of generating NSCs iPSC) was carried out in accordance with the methodology detailed in a previous publication [30].

### Cerebral organoid generation

To generate cortical organoids from iPSCs, we used the protocol described previously [31]. Briefly, feeder-free iPSCs were fed daily with E8 medium for at least 7 days before differentiation. Colonies were dissociated using Accutase (Life Technologies) in PBS (1:1) for 10 min at 37 °C and centrifuged for 3 min at 300 × g. A total of 9,000 live cells were then seeded to 96-well ultra-low attachment tissue culture plates (Thermo Fisher Scientific) in 150 μl neural induction media and kept in suspension under rotation (95 rpm) in the presence of 50 μM ROCK inhibitor (Stem cell Technologies) and kept in suspension under rotation (85 rpm) for 24 h to form embryoid body (EB). To minimize un-directed differentiation, dual SMAD inhibition and canonical WNT inhibition are adopted during this period. On day 2, half of the media was replaced by human neural induction media in the presence of 50 μM ROCK inhibitor to each well. On days 4, 6, and 8, 100 μl medium was placed with 150 μl neural induction without ROCK inhibitor. After 10 days, the organoids were transferred into 6-well ultra-low attachment tissue culture plates (Life Sciences) in neural differentiation media minus vitamin A for the next 8 days using an orbital shaker to induce cerebral organoids. On day 18, the organoids were subsequently matured in neural differentiation media with vitamin A with media changes every 3-4 days. BDNF and ascorbic acid were supplemented to facilitate long-term neural maturation.

### NR treatment of cortical organoids

NR was kindly provided by Evandro Fei Fang, University of Oslo, Norway. One-month-old organoids were treated with 1 mM NR for 2 months. NR was added to the medium and replaced every three days.

### Snap-freezing and embedding

Each human cerebral organoid was fixed in 4% paraformaldehyde in PBS overnight at 4°C, dehydrated with 30% sucrose in phosphate-buffered saline (PBS). The samples were then embedded int gelatin solution (Sigma-Aldrich) and snap frozen in nitrogen. The samples were then embedded in O.C.T. compound (Thermo Fisher Scientific). Cryostat sections (15 µm) were cut and mounted onto slides (Thermo Fisher Scientific).

### Immunofluorescence staining

Mounted sections were incubated for 1 h at room temperature with and blocked using 1 X PBS supplemented with 10% normal goat serum or 5% Bovine Serum Albumin (BSA) and 0.1% Triton X-100 and then incubated with primary antibodies diluted in blocking solution overnight at 4°C. The following primary antibodies were used for immunostaining: Oct4 (1:100; Abcam, cat. no. ab19857), SSEA4 (Abcam, cat. no. ab16287, 1:200), Nestin (1:50, Santa Cruz Biotechnology, cat. no. sc23927), NeuN (1:500; Cell Signaling, cat. no. 24307), MAP2 (1:1000; Abcam, cat. no. ab5392), Tuj1 (1:1000; Abcam, cat. no. ab78078), SOX2 (1:100; Abcam, cat. no. ab97959), NDUFB10 (1:350; Abcam 196019), GFAP (1:500; Abcam cat. no. ab4674), SATB2 (rabbit, 1:400; Abcam cat. no. ab4674), CTIP2 (rat, 1:500; Abcam cat. no. ab18465). Alexa Fluor Dyes (Life Technologies) were used at 1:800 dilution as secondary antibodies. Slides were mounted using ProLong^TM^ diamond antifade mounting medium with DAPI (Life Technologies) and analyzed using the Leica TCS SP8 confocal microscope (Leica Microsystems) or Dragonfly Confocal Microscope (Andor).

### Quantification and image analysis

Image analysis and fluorescent signal quantification were performed with ImageJ software. For staining of GFAP, MAP2, Tuj1 and NDUFB10, the mean fluorescent intensity of each representative field was calculated and compared between groups. For staining of NeuN, SOX2, CTIP2 and SATB2, the number of positive stained nuclei in each representative field was calculated and compared between groups.

### DNA sequencing for *POLG* mutations

Forward and backward oligonucleotide primers were used to amplify the 7 exons and 10 exons of the *POLG* gene. Automated nucleotide sequencing was performed using the Applied Biosystems™ BigDye^®^Terminator v3.1 Cycle Sequencing Kit (Life Technologies, cat. no. 4337454) and analyzed on an ABI3730 Genetic Analyzer with sequencing analyzer software Chromas^Pro^ (Technelysium Pty Ltd, Australia). DNA Chromatogram was aligned with the best matching human sequences in NCBI Trace.

### Transmission electron microscopy (TEM)

Cells were fixed with 4% glutaraldehyde and postfixed with 1% OsO4 in 0.1 mol/L cacodylate buffer containing 0.1% CaCl_2_ at 4°C for 2 hours. Samples were stained with 1% Millipore-filtered uranyl acetate, dehydrated in increasing concentrations of ethanol and infiltrated and embedded in epoxy resin. Ultrathin sections were cut and stained with uranyl acetate and lead citrate. Electron photomicrographs were obtained using a transmission electron microscope (JEM-1230, JEOL).

### ROS measurement

ROS production was measured by flow cytometry. Mitochondrial mass-related intracellular ROS was detected using 30 μM DCFDA (Abcam) and 150 nM MTDR (Life Technologies). Mitochondrial ROS production in relation to mitochondrial mass was detected using 10 μM MitoSOX Red Mitochondrial Superoxide Indicator (Life Technologies) and 150 MTG (Life Technologies). double staining. After staining, stained cells were detached using TrypLE^TM^ Express enzyme (Life Technologies) and neutralized with a medium containing 10% FBS, and then analyzed on a FACS BD Accuri^TM^ C6 flow cytometer (BD Biosciences, San Jose, CA, USA). At least 40,000 events were recorded per sample, and doublets or dead cells were excluded. The results were analyzed using Accuri^TM^ C6 software.

### MtDNA copy number measurement

MtDNA copy number was quantified using flow cytometry and qPCR. For flow cytometry, cells were stained with TFAM and TOMM20. Cells were detached with TrypLE^TM^ Express enzyme and fixed with 1.6% (v/v) PFA (VWR) for 10 min at room temperature. Cells were permeabilized with ice-cold 90% methanol for 20 min at - 20°C after washing with PBS, followed by blocking buffer containing PBS, 0.3M glycine, 5% goat serum, and 1% BSA. closed. Cells were incubated with anti-TFAM antibody conjugated to Alexa Fluor^®^ 488 (Abcam) at 1:400 dilution and anti-TOMM20 antibody conjugated to Alexa Fluor^®^ 488 (Santa Cruz Biotechnology) at a 1:400 dilution. Cells were then analyzed on a BD Accuri^TM^ C6 flow cytometer. At least 40,000 events were recorded per sample, and doublets or dead cells were excluded. The data analysis was performed using Accuri^TM^ C6 software.

For qPCR, DNA was extracted using the QIAGEN DNeasy Blood and Tissue Kit (QIAGEN) according to the manufacturer’s protocol and ND1/APP was measured as previously published method for relative quantification of mtDNA copy number [7].

### Mitochondrial respiratory chain complex measurement

Protein levels of mitochondrial CI, CII, and CIV were accessed using flow cytometry. Cells were detached with TrypLE^TM^ enzyme and were fixed with 1.6% (v/v) PFA (VWR) for 10 minutes at RT, before permeabilized using ice-cold 90% methanol at −20°C for 20 min. The cells were blocked using the blocking buffer mentioned above. Cells were stained with primary antibodies at dilutions 1:1000. The primary antibodies include anti-NDUFB10 (Abcam), anti-SDHA (Abcam) and anti-COX IV (Abcam). The secondary antibodies Alexa Flour^®^ goat Anti-rabbit 488 or Anti-mouse 488 (Thermo Fisher Scientific) were subsequently incubated at dilutions 1:400. The cells were analyzed on BD Accuri^TM^ C6 flow cytometer and data analysis was performed using Accuri^TM^ C6 software. At least 40,000 events were recorded for each sample, doublets or dead cells were excluded.

### MMP measurement

MMPs were measured relative to mitochondrial volume using flow cytometry. Cells were double stained with 100 nM TMRE (Abcam) and 150 nM MTG (Life Technologies) for 45 min at 37 °C. Cells treated with 100 μM Carbonyl cyanide-p-trifluoromethoxyphenylhydrazone (FCCP) (Abcam) were used as a negative control. After washing with PBS, cells were detached with TrypLE^TM^ enzyme and neutralized with medium supplemented with 10% FBS. Cells were then analyzed on a FACS BD Accuri^TM^ C6 flow cytometer. Data analysis was performed using Accuri™ C6 software. At least 40,000 events were recorded per sample, and doublets or dead cells were excluded.

### ATP production

Intracellular ATP production was detected using the Luminescent ATP Detection Assay Kit (Abcam). Cells were grown to 90% confluency in 96-well plates (Life Sciences, cat. no. 3601). ATP measurements were performed according to the manufacturer’s protocol. After cells were lysed, luciferase and luciferin were added, and the emitted light corresponding to the amount of ATP was measured in a Victor^®^ XLight Multimode Plate Reader (PerkinElmer). For each sample, measure 3-6 replicates. To normalize cell numbers for ATP production, cells grown in the same 96-well plate were stained using the Janus Green Cell Normalization Staining Kit (Abcam). OD values at 595 nm were measured by a Labsystems Multiskan Bichromatic plate reader (Titertek Instruments, USA).

### L-lactate measurement

L-lactate production was analyzed by a colorimetric L-lactate assay kit (Abcam) according to the manufacturer’s instructions. Determine the endpoint lactate concentration in a 96-well plate by measuring the initial rate (2 min) of lactate dehydrogenase equilibration between NAD^+^ and NADH. Immediately after the extracellular flux assay, plates were measured at OD 450 nm in a microplate reader (VICTOR™ XLight, PerkinElmer).

### RNA sequencing analysis

Total RNA was isolated from NSCs and iPSCs using RNeasy Mini Kit (QIAGEN). RNA quality and concentration were checked using Bioanalyzer and Qubit^TM^. RNA-seq libraries were established by the PolyA enrichment method. The libraries were sequenced at a depth of over 22 million raw reads, ensuring comprehensive coverage of the transcriptome and sufficient depth for accurate and reliable analysis of gene expression. For iPSCs and NSCs, RNA sequencing analysis was performed by HudsonAlpha. For brain organoid, the RNA sequencing analysis was performed by BGI. FASTQ files were trimmed using Trimmomatic version 0.39 to remove potential Illumina adapters and low quality bases with the following parameters: ILLUMINACLIP: truseq.fa:2:30:10 LEADING:3 TRAILING:3 SLIDINGWINDOW:4:15 [50]. FASTQ files were assessed using fastQC version 0.11.8 prior and following trimming [51]. We used Salmon version 1.0.0 to quantify the abundance at the transcript level with the fragment-level GC bias correction option (*gcBias*flag) and the appropriate option for the library type (*l*flag set to A) against the GENCODE release 32 of the human transcriptome (GRCh38.p13) [52]. Transcript-level quantification was imported into R and collapsed onto gene-level quantification using the tximport R package version 1.8.0 according to the gene definitions provided by the same GENCODE release [53]. The genes in non-canonical chromosomes and scaffolds were filtered out. Genes were filtered out if their level of expression was below 10 reads in more than 75% of the samples based on CPM (count per million). Transcripts counts were aggregated into gene counts. Sample correlations were calculated with the information of library size, cell type and mutation, and sample outliers were excluded. Differential expression analysis was conducted using DEseq2 [54]. Multiple comparisons were adjusted by using the false discovery rate method. Adjusted P value (q value) <0.05 was considered as statistical significance. KEGG pathway and GO enrichment analysis were conducted using Clusterprofiler [55]. To ensure the robustness and reliability of our results, we included three replicates for each clone in the iPSC and NSC samples. Furthermore, for the organoid samples, we included three replicates for the patient lines and four replicates for the control line. In each replicate, we analyzed 4-6 organoids for RNA sequencing, allowing us to capture the inherent biological variability within the samples.

### Statistical analysis

Data were presented as mean ± standard deviation (SD) for the number of samples (n ≥ 3). Distributions were tested for normality using the Shapiro-Wilk test. Outliers were detected using the ROUT method. The Mann-Whitney U-test was used to assess statistical significance for variables with non-normal distribution, while a two-sided Student’s t-test was applied for normally distributed variables. Data were analyzed and figures were produced with GraphPad Prism 8.0.2 software GraphPad Software, Inc). P ≤ 0.05 was considered significant.

## Data availability

The datasets generated and analyzed during the study are included with the Supplemental Information. The RNA sequencing analysis read count data can be accessed in NCBI Gene Expression Omnibus (GEO) data deposit system with an accession number GSE207007. All other data are available from the corresponding author upon request.

## Author’s contributions

K.L contribute to the conceptualization; K.L and Y. H contribute to the methodology; K.L, Z. Y, A.C, B.C.L contribute to the investigation; K.L, Y. H and Z. Z contribute to the writing original draft; all authors contribute to writing review and editing; K.L, L.A.B, E.F.F and G.J.S contribute to the funding acquisition and to the resources; K.L contributes to the supervision. All authors agree to the authorships.

## Acknowledgements

We thank members of the Molecular Imaging Centre and Flow Cytometry Core Facility for their expertise and assistance in confocal imaging and flow cytometry data recording.

---

**Table S1:** The KEGG pathways of the down- and up-regulated DEGs in iPSCs from patient organoids versus controls.

| ID | pathway | pvalue | geneID |
| --- | --- | --- | --- |
| *iPSC downregulated pathways* |  |  |  |
| M00142 | NADH:ubiquinone oxidoreductase, mitochondria | 0.000729 | ND1/ND4/ND5/ND4L |
| M00140 | C1-unit interconversion, prokaryotes | 0.00512 | SHMT2/SHMT1 |
| M00346 | Formaldehyde assimilation, serine pathway | 0.015401 | ENO3/SHMT2/SHMT1 |
| M00158 | F-type ATPase, eukaryotes | 0.028616 | ATP5PO/ATP5PB/ATP5MC3/ATP8 |
| *iPSC upregulated pathways* |  |  |  |
| M00089 | Triacylglycerol biosynthesis | 0.001815 | AGPAT1/LPIN2/AGPAT4/GPAT3/DGAT2/LCLAT1 |
| M00345 | Formaldehyde assimilation, ribulose monophosphate pathway | 0.01069 | PFKM/PFKP |
| M00001 | Glycolysis (Embden-Meyerhof pathway), glucose => pyruvate | 0.017689 | PGAM2/GCK/PFKM/PFKP/PKLR |
| M00056 | O-glycan biosynthesis, mucin type core | 0.024055 | GALNT2/GALNT14/GCNT4/C1GALT1C1L/GALNT18 |

**Table S2:**
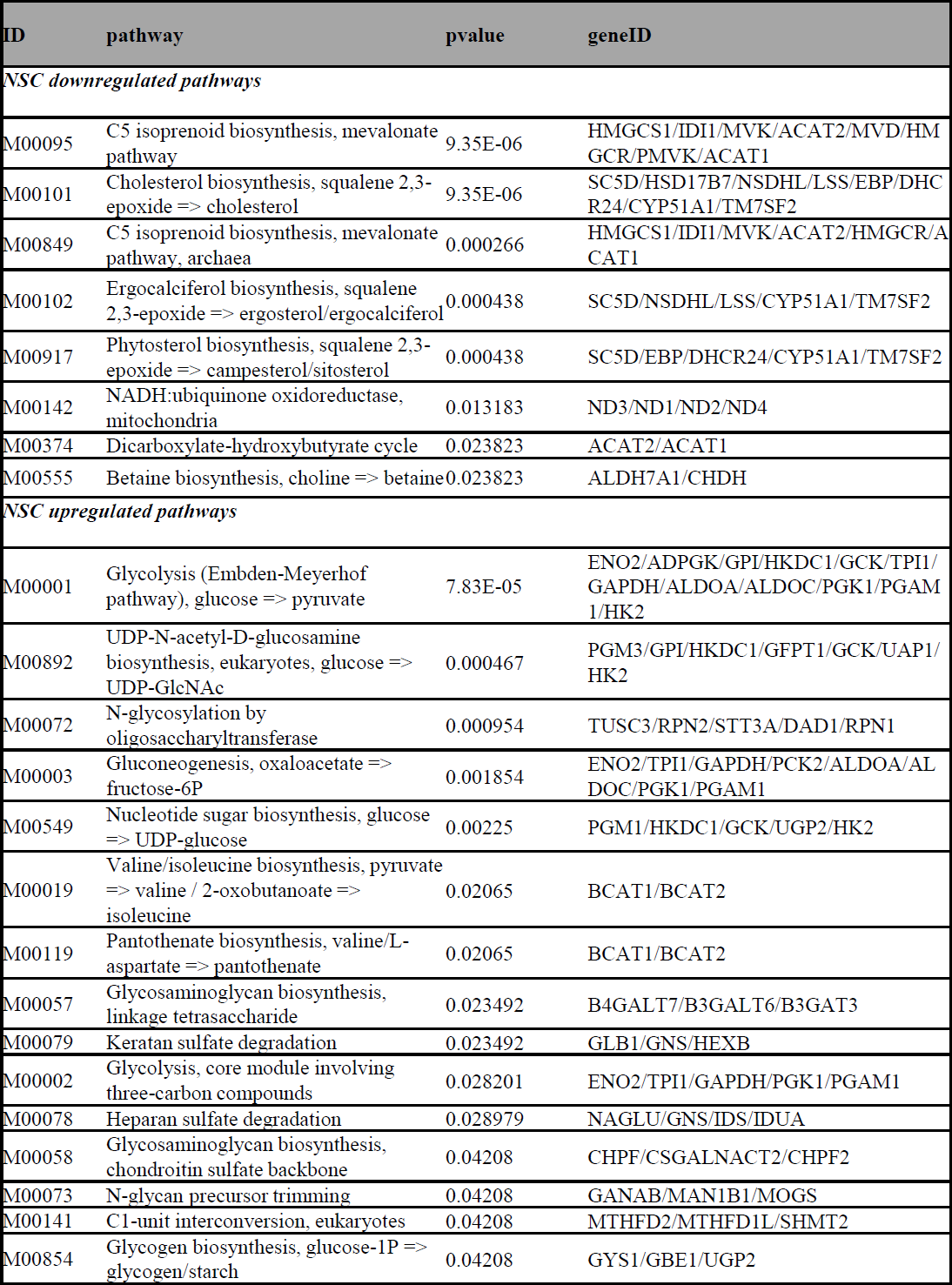
The KEGG pathways of the down- and up-regulated DEGs in NSCs from patient organoids versus controls.

**Table S3:** The mitochondrial related pathways of upregulated DEGs in cortical organoids from patient organoids versus controls.

| ID | pathway | pvalue | core_enrichment |
| --- | --- | --- | --- |
| GO:0019896 | axonal transport of mitochondrion | 0.000186 | TRAK1/AGBL4/FEZ1/SPAST/OPA1/ACTR10/AGTPBP1/UCHL1/MAPT/HAP1/SYBU/NEFL |
| GO:0034643 | establishment of mitochondrion localization, microtubule-mediated | 0.002215 | MAP1S/TRAK1/RHOT1/AGBL4/FEZ1/SPAST/OPA1/ACTR10/AGTPBP1/MAP1B/UCHL1/MAPT/HAP1/SYBU/NEFL |
| GO:0047497 | mitochondrion transport along microtubule | 0.002215 | MAP1S/TRAK1/RHOT1/AGBL4/FEZ1/SPAST/OPA1/ACTR10/AGTPBP1/MAP1B/UCHL1/MAPT/HAP1/SYBU/NEFL |
| GO:0051654 | establishment of mitochondrion localization | 0.007642 | MAP1S/TRAK1/RHOT1/AGBL4/FEZ1/SPAST/OPA1/ACTR10/MARK1/AGTPBP1/MAP1B/UCHL1/MAPT/HAP1/SYBU/NEFL |
| GO:0006120 | mitochondrial electron transport, NADH to ubiquinone | 0.015314 | NDUFV1/NDUFB6/NDUFAB1/NDUFS5/NDUFS7/NDUFB2/NDUFS1/NDUFAF1/ND3/NDUFA6/NDUFC2-KCTD14/ND2/NDUFA5/ND1/PINK1/NDUFA9/ND5/ND4L/ND4/SNCA |
| GO:0090646 | mitochondrial tRNA processing | 0.034206 | TRMT61B/TRIT1/TRMT5/MTO1/RPUSD4/HSD17B10/PRORP/TRNT1/PUS1/ELAC2/CDK5RAP1 |
| GO:0000963 | mitochondrial RNA processing | 0.036468 | FASTKD2/FASTKD3/TRMT61B/TRIT1/SUPV3L1/TBRG4/TRMT5/MTO1/RPUSD4/HSD17B10/PRORP/TRNT1/PUS1/FASTKD5/ELAC2/FASTKD1/CDK5RAP1 |
| GO:0005747 | mitochondrial respiratory chain complex I | 0.040214 | NDUFA10/NDUFB3/NDUFA8/NDUFA4/NDUFV1/NDUFB6/NDUFAB1/NDUFS5/NDUFA12/NDUFS7/NDUFB2/NDUFS1/NDUFAF1/ND3/FOXRED1/NDUFA6/NDUFC2-KCTD14/ND2/NDUFA5/ND1/NDUFA9/ND5/ND4L/ND4 |
| GO:0032981 | mitochondrial respiratory chain complex I assembly | 0.040874 | NDUFAF6/TMEM126B/NDUFB6/NDUFAB1/NDUFS5/NDUFA12/NDUFS7/NDUFB2/NDUFS1/NDUFAF1/FOXRED1/NDUFAF2/NDUFA6/TIMM21/BCS1L/ND2/NDUFA5/ND1/NDUFA9/NDUFAF4/NDUFAF5/ND5/ND4 |

**Table S4:** The axon and synaptic related pathways of the downregulated DEGs in cortical organoids from patient organoids versus controls.

| ID | pathway | pvalue | core_enrichment |
| --- | --- | --- | --- |
| GO:0019896 | axonal transport of mitochondrion | 0.000186 | TRAK1/AGBL4/FEZ1/SPAST/OPA1/ACTR10/AGTPBP1/U<br>CHL1/MAPT/HAP1/SYBU/NEFL |
| GO:0034643 | establishment of mitochondrion localization, microtubule-mediated | 0.002215 | MAP1S/TRAK1/RHOT1/AGBL4/FEZ1/SPAST/OPA1/ACT<br>R10/AGTPBP1/MAP1B/UCHL1/MAPT/HAP1/SYBU/NEFL |
| GO:0047497 | mitochondrion transport along microtubule | 0.002215 | MAP1S/TRAK1/RHOT1/AGBL4/FEZ1/SPAST/OPA1/ACT<br>R10/AGTPBP1/MAP1B/UCHL1/MAPT/HAP1/SYBU/NEFL |
| GO:0051654 | establishment of mitochondrion localization | 0.007642 | MAP1S/TRAK1/RHOT1/AGBL4/FEZ1/SPAST/OPA1/ACT<br>R10/MARK1/AGTPBP1/MAP1B/UCHL1/MAPT/HAP1/SYB<br>U/NEFL |
| GO:0006120 | mitochondrial electron transport, NADH to ubiquinone | 0.015314 | NDUFV1/NDUFB6/NDUFAB1/NDUFS5/NDUFS7/NDUFB<br>2/NDUFS1/NDUFAF1/ND3/NDUFA6/NDUFC2-KCTD14/ND2/NDUFA5/ND1/PINK1/NDUFA9/ND5/ND4L/<br>ND4/SNCA |
| GO:0090646 | mitochondrial tRNA processing | 0.034206 | TRMT61B/TRIT1/TRMT5/MTO1/RPUSD4/HSD17B10/PRO<br>RP/TRNT1/PUS1/ELAC2/CDK5RAP1 |
| GO:0000963 | mitochondrial RNA processing | 0.036468 | FASTKD2/FASTKD3/TRMT61B/TRIT1/SUPV3L1/TBRG4/<br>TRMT5/MTO1/RPUSD4/HSD17B10/PRORP/TRNT1/PUS1/<br>FASTKD5/ELAC2/FASTKD1/CDK5RAP1 |
| GO:0005747 | mitochondrial respiratory chain complex I | 0.040214 | NDUFA10/NDUFB3/NDUFA8/NDUFA4/NDUFV1/NDUFB<br>6/NDUFAB1/NDUFS5/NDUFA12/NDUFS7/NDUFB2/NDU<br>FS1/NDUFAF1/ND3/FOXRED1/NDUFA6/NDUFC2-KCTD14/ND2/NDUFA5/ND1/NDUFA9/ND5/ND4L/ND4 |
| GO:0032981 | mitochondrial respiratory chain complex I assembly | 0.040874 | NDUFAF6/TMEM126B/NDUFB6/NDUFAB1/NDUFS5/ND<br>UFA12/NDUFS7/NDUFB2/NDUFS1/NDUFAF1/FOXRED1/<br>NDUFAF2/NDUFA6/TIMM21/BCS1L/ND2/NDUFA5/ND1/<br>NDUFA9/NDUFAF4/NDUFAF5/ND5/ND4 |

**Table S5:**
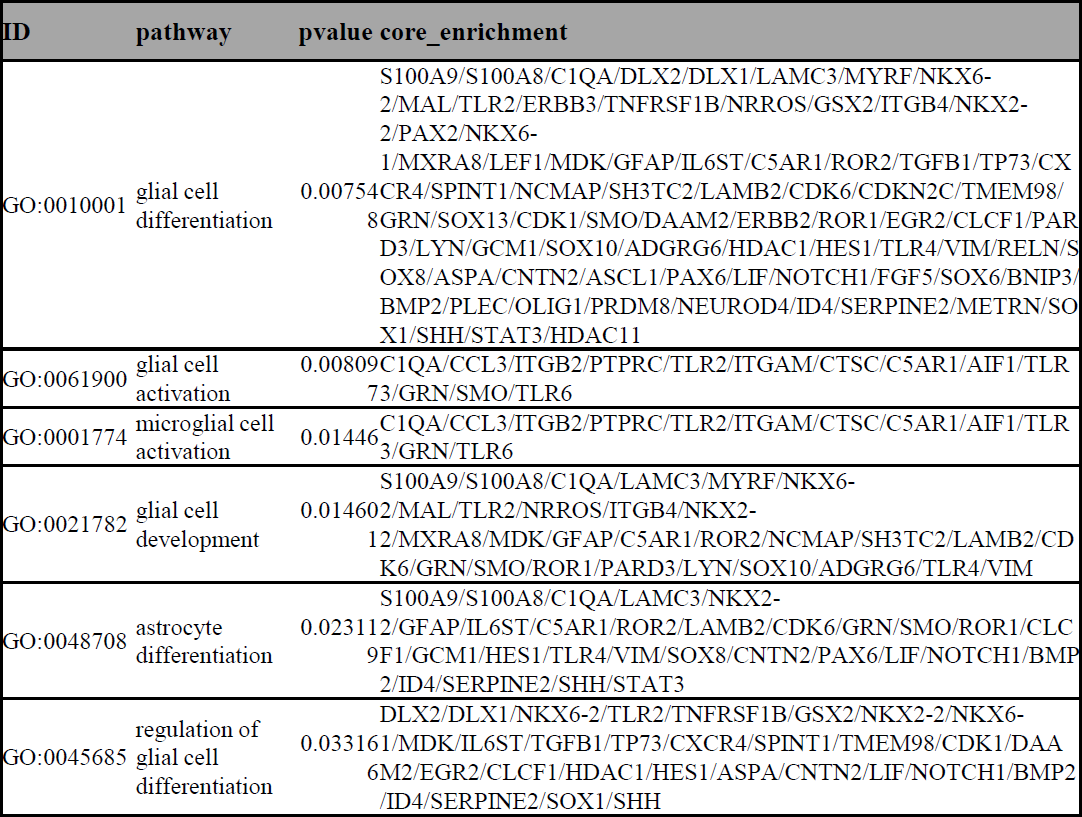
The astrocyte related pathways of the upregulated DEGs in cortical organoids from patient organoids versus controls.

**Table S6:** The neuroinflammation related pathways of the downregulated DEGs in cortical organoids from patient organoids versus controls.

| ID | pathway | pvalue | core_enrichment |
| --- | --- | --- | --- |
| GO:0050727 | regulation of inflammatory response | 1.22E-09 | KRT1/S100A9/PLA2G2A/S100A8/APOA1/CD28/CASP4/CCL3/PLA2G7/CREB3L3/ALOX5AP/GATA3/PTGER4/NR1H4/ACP5/VAMP8/MMP9/PTPRC/HCK/IL1R1/BIRC3/LGALS1/CASP1/SYK/TLR2/FOXF1/SLAMF8/TNFRSF1B/AHSG/C3/ANXA1/PYCARD/LRRC19/ETS1/ISL1/TNFRSF11A/CCN4/NCF1/ALOX5/ADAMTS12/MGST2/GPER1/STING1/GPSM3/CD200R1/SOCS3/DUOXA2/NFKBIZ/MDK/IFI35/IL6ST/CTSC/PTGES/AGTR1/NMI/TEK/HGF/PPARG/TNFAIP3/OSMR/NUPR1/TLR3/ZFP36/PLCG2/SCGB1A1/HLA-E/APOE/PBK/CEBPA/KLF4/NOD2/AGT/LACC1/DUOXA1/TNFRSF1A/AOAH/WNT5A/GRN/METRNL/NLRX1/MYD88/TLR7/SMAD3/MAPK13/LYN/SERPINF1/PTGS2/ADORA2B/PIK3CG/PPARA/TREX1/CEBPB |
| GO:0002526 | acute inflammatory response | 9.16E-06 | SERPINA1/S100A8/APOA2/CD163/CREB3L3/ALOX5AP/SERPINA3/GATA3/FN1/A2M/AHSG/C3/TNFRSF11A/IL6R/PLSCR1/IL6ST/PTGES/ASS1/B4GALT1/F3/OSMR/NUPR1/KL/HLA-E/TACR1/TREM1/ANO6/SERPINF2/PTGS2/PIK3CG/HP/CEBPB/TFRC |
| GO:0050729 | positive regulation of inflammatory response | 2.84E-05 | S100A9/PLA2G2A/S100A8/CD28/CCL3/PLA2G7/CREB3L3/ALOX5AP/PTGER4/VAMP8/LGALS1/CASP1/TLR2/C3/PYCARD/ETS1/TNFRSF11A/CCN4/MGST2/GPSM3/NFKBIZ/MDK/IFI35/IL6ST/CTSC/AGTR1/NMI/OSMR/NUPR1/TLR3/PLCG2/HLA-E/CEBPA/AGT/TNFRSF1A/WNT5A/GRN/TLR7/MAPK13/PTGS2/ADORA2B/PIK3CG/CEBPB |
| GO:0002532 | production of molecular mediator involved in inflammatory response | 0.001498 | H19/ALOX5AP/VAMP8/IL4R/SYK/SLAMF8/PYCARD/NCF1/ALOX5/CD36/GPSM3/CLEC7A/NOD2/SNAP23/TICAM1/ZC3H12A/GRN/APOD/ADCY7/MYD88/TLR6/LYN/PPARA/TLR4/IL17RA/NOS2 |
| GO:0061702 | inflammasome complex | 0.002189 | CASP4/CASP1/PYCARD/GSDMD |
| GO:0002523 | leukocyte migration involved in inflammatory response | 0.002256 | S100A9/S100A8/TRIM55/SLAMF8/ALOX5/MDK |
| GO:0150076 | neuroinflammatory response | 0.003557 | C1QA/CCL3/ITGB2/MMP9/PTPRC/TLR2/TNFRSF1B/ITGAM/CD200R1/CTSC/C5AR1/NUPR1/AIF1/TLR3/PLCG2/GRN/SMO/ITGB1/TLR6/PTGS2 |
| GO:0002269 | leukocyte activation involved in inflammatory response | 0.016619 | C1QA/CCL3/ITGB2/PTPRC/TLR2/ITGAM/CTSC/C5AR1/AIF1/TLR3/GRN/MYD88/TLR6 |
| GO:0002437 | inflammatory response to antigenic stimulus | 0.028382 | CD28/GATA3/HCK/SYK/A2M/CD68/C3/IL5RA/HLA-E/NOD2/HMGB2/LYN/TREX1/NPY/RHBDF2/NOTCH1/NOTCH2/GPX1 |
| GO:0150077 | regulation of neuroinflammatory response | 0.030303 | CCL3/MMP9/PTPRC/TNFRSF1B/CD200R1/CTSC/NUPR1/PLCG2/GRN/PTGS2 |
| GO:0002544 | chronic inflammatory response | 0.032217 | S100A9/S100A8/THBS1/TNFAIP3 |
| GO:0002675 | positive regulation of acute inflammatory response | 0.043678 | CREB3L3/ALOX5AP/C3/TNFRSF11A/IL6ST/OSMR/HLA-E/PTGS2/PIK3CG |

**Table S7:** The mitochondrial related pathways of the upregulated DEGs in cortical organoids from patient organoids with NR treatment versus non-treated patient organoids.

| ID | pathway | pvalue | core_enrichment |
| --- | --- | --- | --- |
| GO:0006123 | mitochondrial electron transport, cytochrome c to oxygen | 0.001645 | MTCO2P12/COX2/COX1/COX3 |
| GO:0005753 | mitochondrial proton-transporting ATP synthase complex | 0.014619 | ATP8/ATP6 |
| GO:0019896 | axonal transport of mitochondrion | 0.022516 | UCHL1/SYBU/HAP1/MAPT/FEZ1/NEFL/SPAST/HSBP1/TRAK1/AGTPBP1/MGARP |
| GO:0034643 | establishment of mitochondrion localization, microtubule-mediated | 0.022935 | UCHL1/SYBU/HAP1/MAPT/FEZ1/NEFL/SPAST/MAP1B/HSBP1/TRAK1/AGTPBP1/WASF1/MGARP |
| GO:0047497 | mitochondrion transport along microtubule | 0.022935 | UCHL1/SYBU/HAP1/MAPT/FEZ1/NEFL/SPAST/MAP1B/HSBP1/TRAK1/AGTPBP1/WASF1/MGARP |
| GO:1904923 | regulation of autophagy of mitochondrion in response to mitochondrial depolarization | 0.034473 | HK2/MFN2/PRKN/PINK1 |
| GO:1904925 | positive regulation of autophagy of mitochondrion in response to mitochondrial depolarization | 0.042801 | HK2/MFN2/PRKN/PINK1 |

**Table S8:** The axon and synaptic related pathways of the downregulated DEGs in cortical organoids from patient organoids with NR treatment versus non-treated patient organoids.

| ID | pathway | pvalue | core_enrichment |
| --- | --- | --- | --- |
| GO:0032228 | regulation of synaptic transmission, GABAergic | 0.007103 | TACR1/SYN3/CLN3/NPAS4/HTR1B/PLCL1/NPY5R/HAP1/ADORA1/PRKCE/NLGN2/CNR1 |
| GO:0098962 | regulation of postsynaptic neurotransmitter receptor activity | 0.007839 | NPTX2/CACNG4/NPTXR/CNIH2/SHISA7/DLGAP3/DLGAP2/DLGAP4/AKAP9/HOMER1 |
| GO:0097091 | synaptic vesicle clustering | 0.012461 | SYN3/SYN1/CDH2/BRSK2/BRSK1/NLGN2/PCLO/RAB3A/SYNDIG1/SYN2 |
| GO:0060080 | inhibitory postsynaptic potential | 0.022009 | NPAS4/NLGN3/GRIK2/NLGN2/CHRNA4/INSYN1/ABAT/RIMS1/UNC13B |
| GO:0048172 | regulation of short-term neuronal synaptic plasticity | 0.028901 | CLN3/SYT4/GRIK2/SHISA7/SYP/RAB3A/SHISA8 |
| GO:0048489 | synaptic vesicle transport | 0.029987 | KIF5A/PRKN/KIF5C/MAP2/AP3M2/AP3B2/CNIH2/LIN7A/LRRK2/PINK1/TRIM46/BLOC1S1/RAB3A/AP3S2/SNCA/AP3M1/SNAP91 |
| GO:0098845 | postsynaptic endosome | 0.033863 | ARC/ZDHHC2/PSD/GRIPAP1/SH3GL3/PICK1/CLSTN1/RAB4A/NSG1/STX12 |
| GO:0097104 | postsynaptic membrane assembly | 0.042748 | NLGN3/NLGN4X/NRXN2/NRXN1/CDH2/NLGN2 |
| GO:0099243 | extrinsic component of synaptic membrane | 0.043173 | CNKSR2/SCRIB/KPNA2/AKAP9/FARP1/SNAP91/RNF10/HIP1 |

**Table S9:**
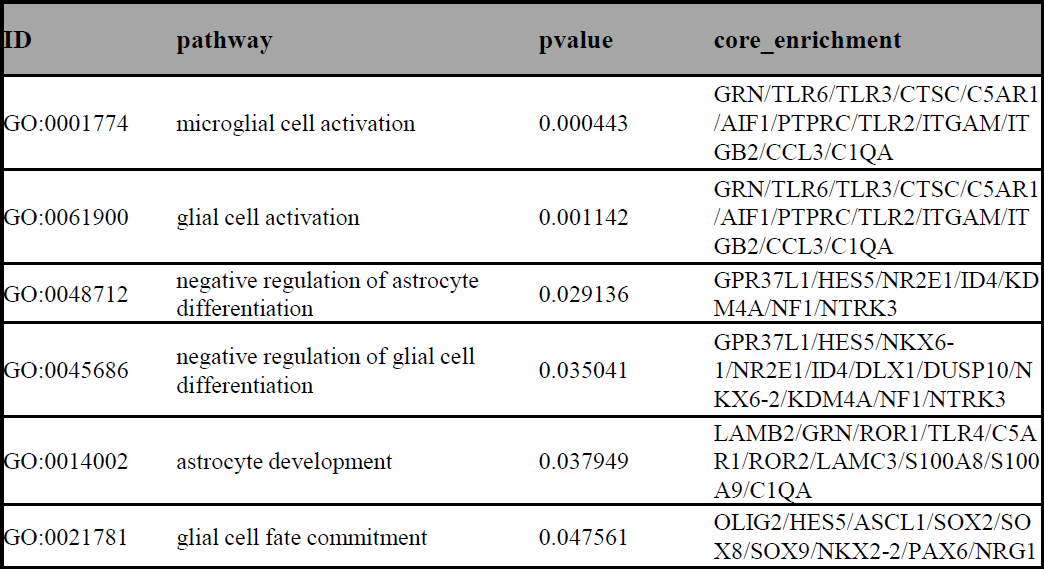
The astrocyte related pathways of the upregulated DEGs in cortical organoids from patient organoids with NR treatment versus non-treated patient organoids.

| ID | pathway | pvalue | core_enrichment |
| --- | --- | --- | --- |
| GO:0001774 | microglial cell activation | 0.000443 | GRN/TLR6/TLR3/CTSC/C5AR1/AIF1/PTPRC/TLR2/ITGAM/ITGB2/CCL3/C1QA |
| GO:0061900 | glial cell activation | 0.001142 | GRN/TLR6/TLR3/CTSC/C5AR1/AIF1/PTPRC/TLR2/ITGAM/ITGB2/CCL3/C1QA |
| GO:0048712 | negative regulation of astrocyte differentiation | 0.029136 | GPR37L1/HES5/NR2E1/ID4/KDM4A/NF1/NTRK3 |
| GO:0045686 | negative regulation of glial cell differentiation | 0.035041 | GPR37L1/HES5/NKX6-1/NR2E1/ID4/DLX1/DUSP10/NKX6-2/KDM4A/NF1/NTRK3 |
| GO:0014002 | astrocyte development | 0.037949 | LAMB2/GRN/ROR1/TLR4/C5AR1/ROR2/LAMC3/S100A8/S100A9/C1QA |
| GO:0021781 | glial cell fate commitment | 0.047561 | OLIG2/HES5/ASCL1/SOX2/SOX8/SOX9/NKX2-2/PAX6/NRG1 |

**Table S10:**
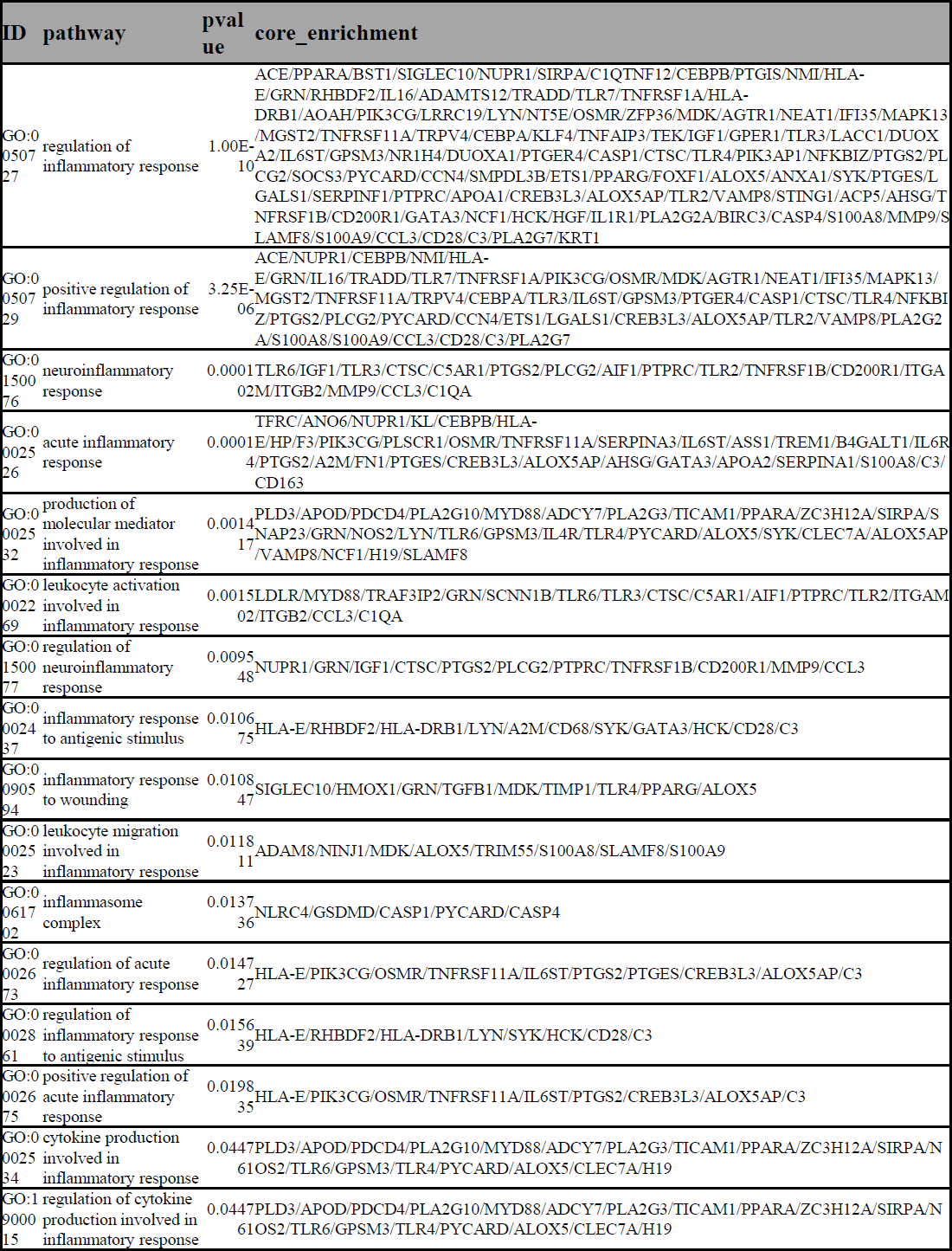
The neuroinflammation related pathways of the upregulated DEGs in cortical organoids from patient organoids with NR treatment versus non-treated patient organoids.

## Supplemental figures

**Supplementary Figure 1:**
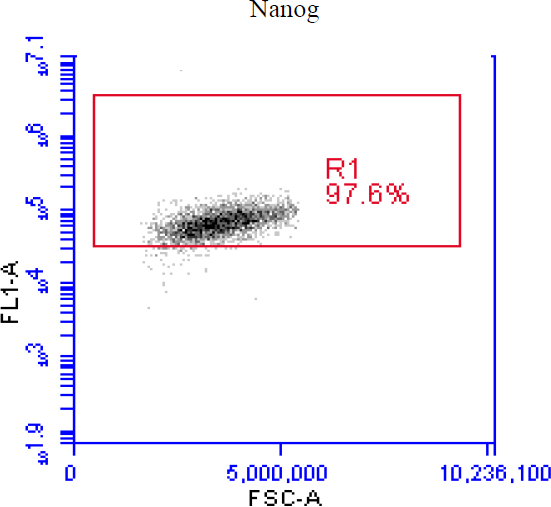
Flow cytometry analysis of Nanog-positive cells.

**Supplementary Figure 2:**
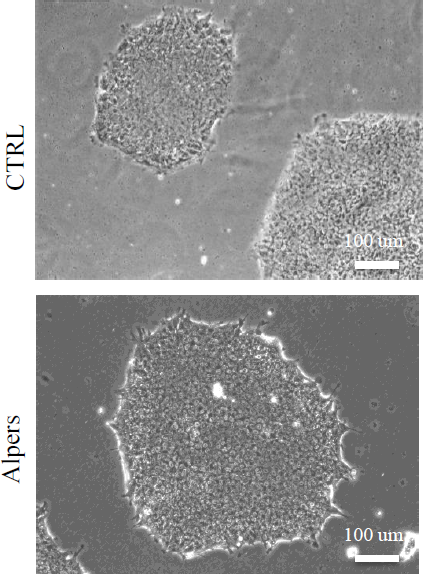
Phase-contrast images in control and Alpers’ iPSCs. Scale bar is 100 µm.

**Supplementary Figure 3:**
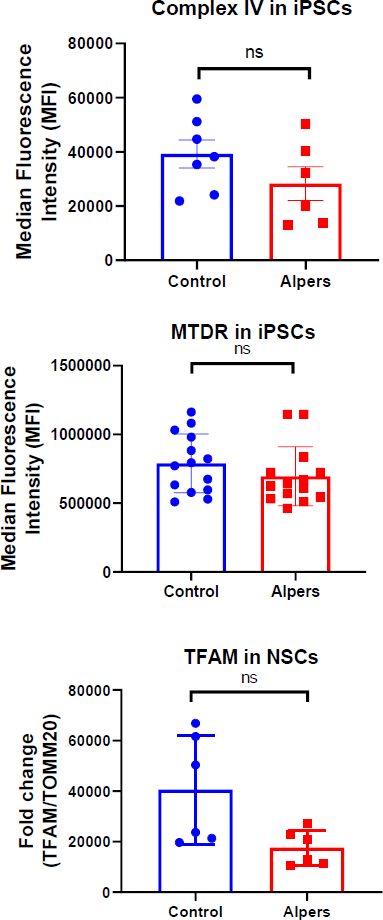
Flow cytometric analysis of CIV levels and MTDR in iPSCs, and TFAM expression in NSCs.

**Supplementary Figure 4:**
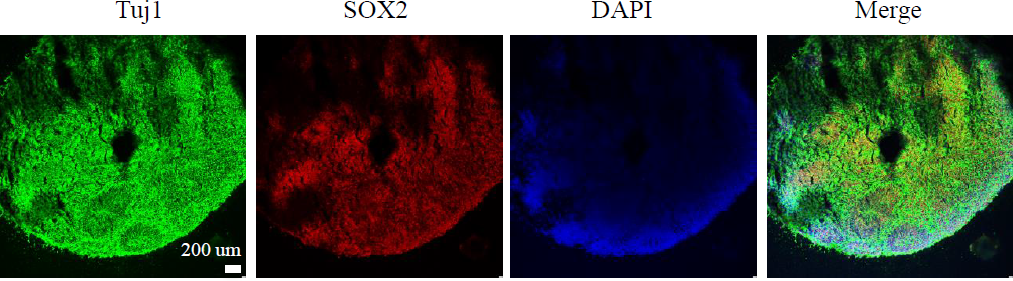
Fluorescent staining of cortical organoid section using neuron marker Tuj1 and neural progenitor marker SOX2 at day 40 in control line. Nuclei are stained with DAPI (blue). Scale bar is 200 µm.

**Supplementary Figure 5:**
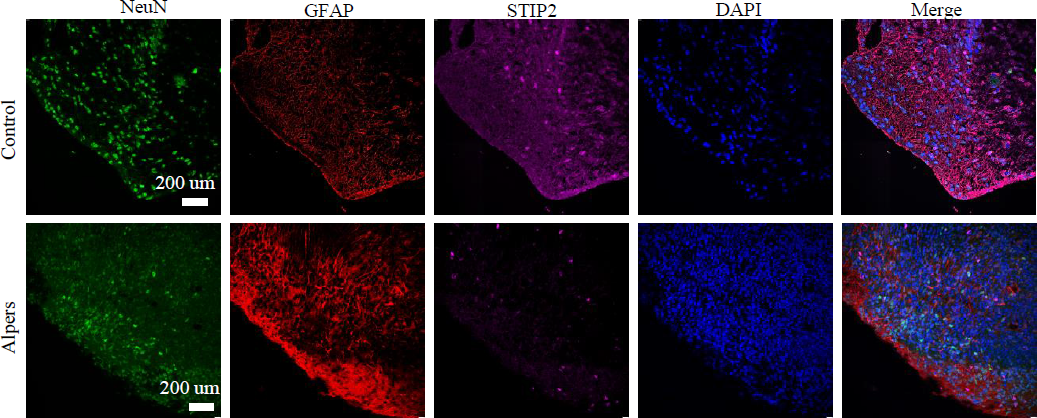
Fluorescent staining of cortical organoid section using neuron marker NeuN, astrocyte marker GFAP, and cortical neuron marker MAP2 at day 30 in control and Alpers’ line. Nuclei are stained with DAPI (blue) Scale bar is 200 µm.

**Supplementary Figure 6:**
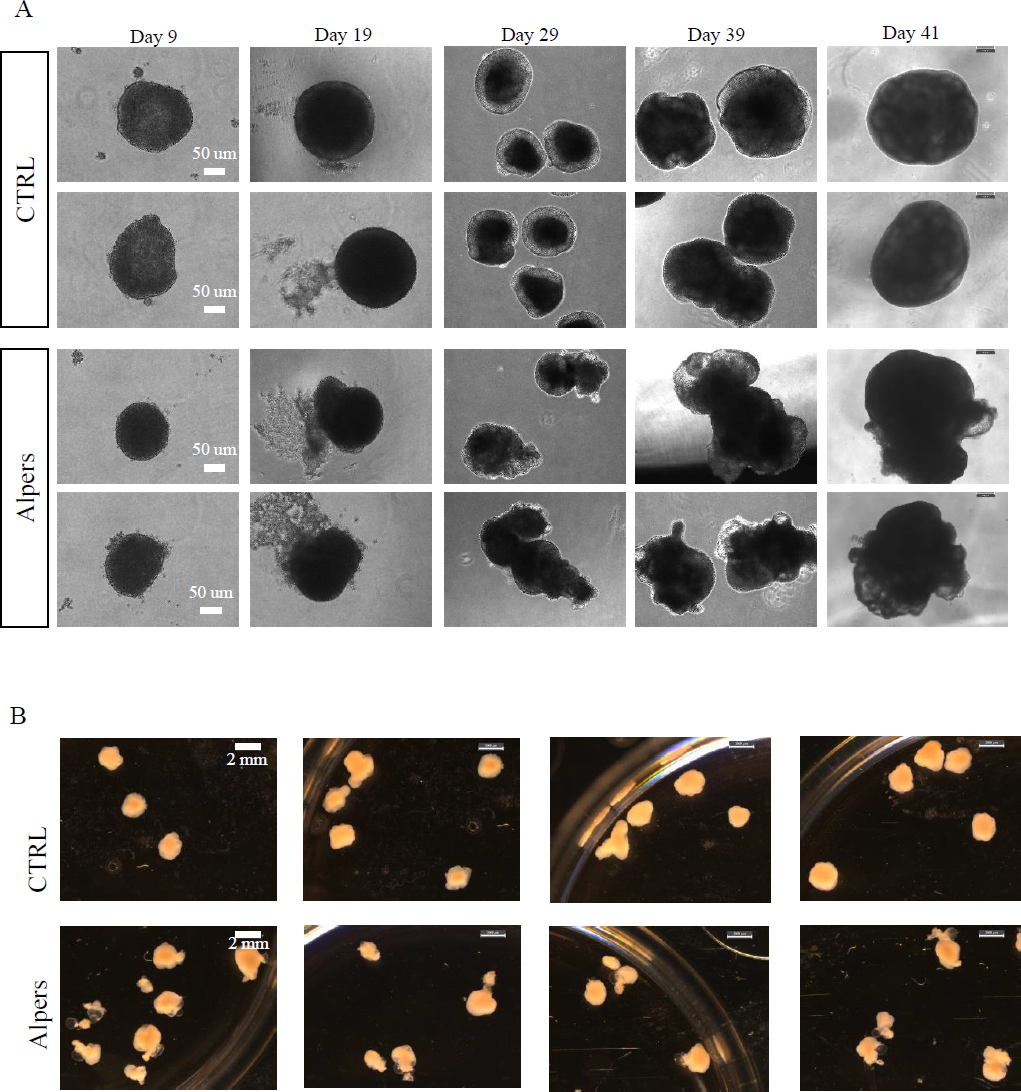
Phase-contrast images of cortical organoids of control and Alpers’ patient line during the differentiation (A) and at day 37 (B) Scale bar is 100 µm or 2 mm.

**Supplementary Figure 7:**
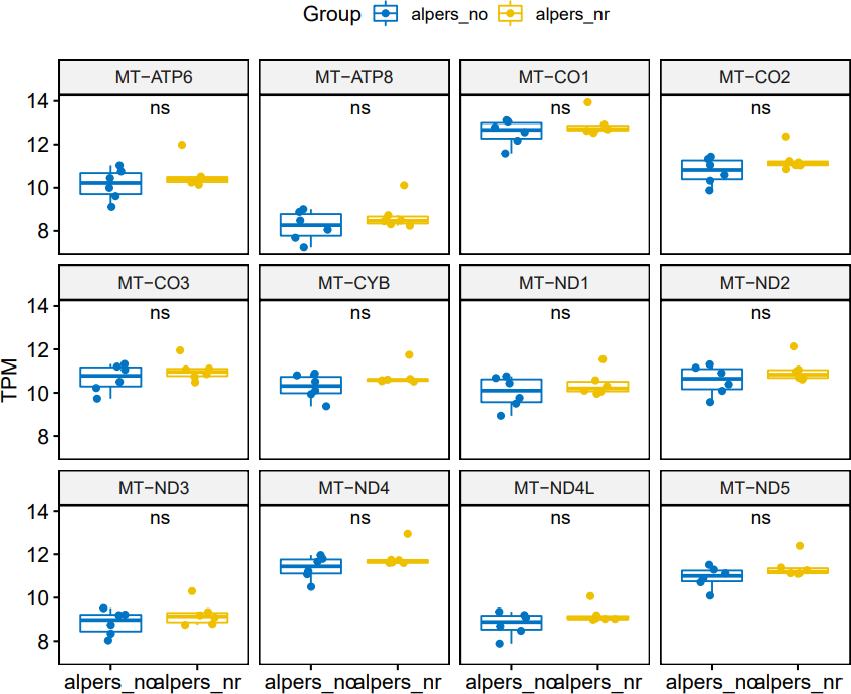
RNA expressions of mitochondrial genes in Alpers’ organoids before and after NR treatment.

